# WispLab: open-source browser workbenches for laboratory image and qPCR analysis with reference-based verification

**DOI:** 10.64898/2026.10.18.759898

**Authors:** Yuan Gao, Sumin Sun

**Affiliations:** Department of Oncology, The Affiliated Huaian No.1 People’s Hospital of Nanjing Medical University, Huai’an, China; Nanjing Normal University, Nanjing, China

**Author notes:** Corresponding authors:* Yuan Gao,; Sumin Sun. These authors contributed equally to this work and share first authorship.

## Abstract

Western blot densitometry, histological image quantification and relative qPCR analysis require choices about regions, backgrounds, thresholds, normalization and statistical comparisons. WispLab connects these operations in three open-source browser workbenches with Chinese and English interfaces, visible parameters and downloadable results. Routine quantitative processing runs locally; an optional external vision service in the Western blot tool is a separate data-transfer pathway. We assessed selected operations using separately implemented references and constructed inputs. For four public Western blot panels containing 21 measured lanes, local-background net signals and reference-lane ratios matched a Python reference at the recorded precision. A six-lane synthetic 16-bit design, duplicated in two files, yielded identical net signals in 12 measurements. Histology verification distinguished successful processing of 35 images from an H-DAB area comparison on eight images and ten regions. After correcting region matching and aligning preprocessing in the reference, positive-area fractions differed by at most 0.152 percentage points. qPCR verification used a public 60-well export and a constructed example; a fresh run of the fixed source version passed 422 specified checks after correction of a deep-tail defect in Dunnett-adjusted probabilities. These tests establish numerical agreement for specified inputs and methods, rather than biological accuracy, clinical validity or superiority over existing software. Source archives, comparison records and reproducible plotting scripts document the supported claims and the remaining validation gaps.

## Introduction

Routine laboratory quantification combines image measurement or instrument exports with background correction, normalization and statistical analysis. Established software includes ImageJ and Fiji for general image analysis [1,2], CellProfiler for high-throughput pipelines [3], QuPath for digital pathology [4], IHC Profiler for ImageJ-based immunohistochemistry scoring [5] and dedicated qPCR tools such as Auto-qPCR [6]; the wider landscape of biological-imaging software has been reviewed elsewhere [7]. The practical problem addressed here is to connect a defined set of operations with inspectable parameters and a direct route from input to exported results.

For Western blots, measurement regions and background subtraction affect densitometric readouts and need to be documented [8], and defined experimental designs and analysis practices have been proposed to improve quantitative reliability [9-11]. Dedicated software also targets Western blot design and analysis, such as the blotRig application for counterbalanced layouts and mixed-model statistics [12]. In immunohistochemistry (IHC), colour deconvolution separates stain contributions [13], thresholding and region selection determine positive-area fractions, and weighted semiquantitative scores such as the H-score and the immunoreactive score remain in routine use [14,15]. Relative qPCR analysis adds reference-gene selection and amplification-efficiency assumptions [16] to the ΔΔCt or Pfaffl calculation [17,18], and available software for these steps has been surveyed [19]. MIQE and its updated guidance describe the experimental and reporting information needed to interpret qPCR results [20,21]. Providing calculations does not by itself establish compliance with those guidelines.

WispLab comprises blot-workbench for Western blots, pathology-workbench for IHC, routine and special stains and fluorescence-image readouts, and pcr-workbench for relative qPCR analysis. The tools use established methods; the contribution is their implementation in linked browser workflows with bilingual interfaces, version tracking and explicit numerical verification. Browser access is not unique to WispLab: Auto-qPCR already provides a web interface [6]. No comparative usability or performance advantage is claimed.

This study asks whether selected WispLab calculations agree with separately implemented references on documented inputs, and whether the evidence can be traced to measurement settings and source versions. We report numerical comparisons, failures in both application and verification code, and the boundaries of the resulting evidence.

## Implementation

### Architecture and data handling

Each workbench is implemented in TypeScript with a React interface and a Vite build. Numerical routines are separated from interface components, allowing selected functions to be called by a scripted harness. Routine calculations run in the browser, with projects and results stored or exported locally (Figure 1). Static hosting serves application files and does not perform these routine calculations.

**Figure 1.**
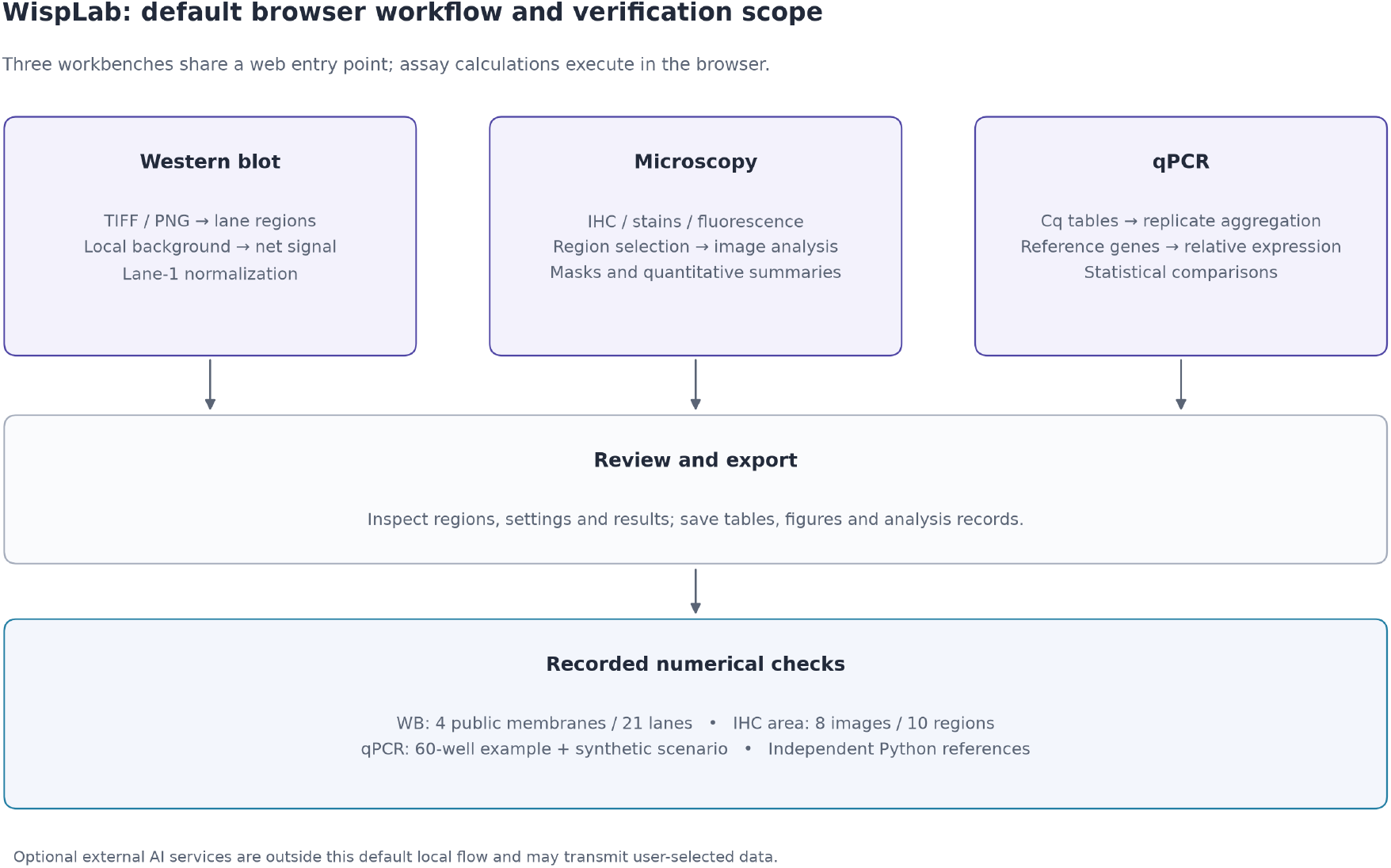
Workflow and verification scope. The three browser workbenches accept Western-blot images, microscopy images or qPCR Cq tables. Users inspect settings and regions before exporting results. Default computations run in the browser; the optional external vision service is a separate data-transfer pathway. The lower panel summarizes the numerical comparisons: four WB panels with 21 measured lanes, eight H-DAB images with ten regions, and a public 60-well qPCR example supplemented by a constructed scenario. These are implementation checks against separately written Python references, not clinical validation or comparisons of biological accuracy.

Local computation is a workflow property, not a blanket promise that no data can leave the device. The Western blot tool has an optional bring-your-own-key vision integration. When the user submits a vision request, it sends an image to the endpoint configured by the user. This service was not needed for the comparisons reported here. The pathology tool supports a user-supplied ONNX segmentation model; that optional pathway was not independently validated.

Each tool has one source repository. A coordinated release process builds for GitHub Pages and the corresponding WispLab domain, and a machine-readable release manifest identifies the source revision of each deployment. Live applications can change. Fixed tags and version-specific archives identify the research releases; the website is not the sole record of a studied version.

### Western blot module

The Western blot tool measures decoded raster pixel values rather than the rendered preview, including a supported unsigned 16-bit TIFF path. Users define band and background rectangles manually or inspect proposals generated from image profiles. Automatic proposals require review and are not treated as validated detection of every band in an arbitrary membrane.

For a band rectangle containing A pixels, pixel sum S and adjacent-background mean B, local-background net signal is **AB − S for dark bands** and **S − AB for light bands**. The implementation retains the subtraction’s sign; it does not take an absolute value. The tool also offers rolling-ball background subtraction and normalization to a selected reference lane or matched loading-control measurement. The real-image comparison reported here used local background and a within-panel reference lane. It does not validate every background or normalization option.

Exports expose measurement regions, results and settings. The reference used the same rectangles and signal convention, testing decoding and measurement agreement conditional on region selection. This is distinct from benchmarking automatic lane localization.

### IHC, staining and fluorescence module

The interface separates H-DAB IHC, routine or special staining, and fluorescence workflows. H&E and trichrome presets are available within the staining workflow. Stain separation uses a matrix-inversion colour-deconvolution formulation [13]. For 8-bit intensity I, optical density is calculated as OD = clip[0,255]{255 ln[255/(I + 1)]/ln(255)}, with endpoint clipping before deconvolution. Normalized stain vectors and a residual vector define the stain matrix. In the evaluated H-DAB path, negative extracted channel values are clipped to zero, a fifth-percentile background estimate is obtained from a 256-bin histogram and subtracted, and a threshold is selected from the corrected channel.

Positive area is the fraction of pixels whose corrected value is strictly greater than the threshold. The evaluated procedure uses Otsu threshold selection [22] for the entire image and applies it to each region of interest (ROI). Selecting a new threshold in each ROI would test a different procedure. Comparisons retained both independently selected thresholds and a reference channel evaluated at the application’s threshold.

Other outputs include optical-density summaries, nuclear counts, labeling index and a nucleus-based H-score implementation [14]. The latter assigns weights of 0, 1, 2 or 3 according to mean intranuclear signal, dividing the interval from the positivity threshold to 255 into equal thirds, and reports 100 times the mean weight. This is not a validated membrane or cytoplasmic clinical H-score. Nuclear counts, H-score and physical-density estimates were not independently validated by the area comparison.

The fluorescence workflow provides channel-based statistics, including Pearson and Manders-type readouts [23,24]. Processing an RGB composite exercises the interface but does not validate colocalization from raw acquisition channels. Background, thresholds and channel selection remain part of interpretation; image-channel association does not establish molecular interaction.

### qPCR module

The qPCR workflow parses supported instrument-export layouts, maps sample and target columns, groups technical replicates and calculates reference-normalized expression. It offers ΔΔCt/Livak and efficiency-corrected Pfaffl calculations [17,18], with selected reference genes and a specified control group. Multiple reference genes are pooled in the Cq domain. Replicate-level and sample-level summaries remain distinct in the analysis.

Inferential routines include Welch comparisons on ΔCt [25] and one-way analysis of variance followed by control-versus-treatment comparisons with a Dunnett adjustment [26]. No-template-control and melting-curve summaries are available as quality-control aids but were not independently validated here. Technical replicate wells were used to exercise numerical routines; their number must not be interpreted as the number of independent biological replicates. The p-values below verify computations and do not support a new biological conclusion about the source dataset.

## Verification methods

### Evidence scope and source versions

We distinguished functional execution, reference comparisons of specified measurements, checks against constructed inputs, and regression tests. Their denominators are not interchangeable (Table 1). Historical records were retained, while subsequent audits and reruns were written separately.

The paper-release tag is `wisplab-paper-2026-10-17` in each repository. Historical notes identify intended revisions, but the original result JSON files do not all embed an execution-time Git SHA. A tag listed beside an old result therefore does not prove which revision ran. On 18 October 2026, we extracted fixed source revisions with Git and reran the PCR pipeline, real-image and synthetic WB measurements, and all 35 histology inputs. Source and input hashes, commands and software versions were recorded. The runs reused existing dependency installations, whose versions were recorded; they were not clean dependency installations. The archived historical records remain available for the correction audit and are not used as substitutes for the newly recorded execution provenance.

Reference implementations were written separately from the application code. They were not assumed infallible: matrix layout, image orientation, region matching and preprocessing definitions were audited. Agreement between implementations can expose inconsistencies, but cannot exclude shared errors or establish the biological suitability of a method.

### Western blot inputs

The real-image comparison used four public vendor panels: β-actin, phosphorylated ERK1/2, phosphorylated AKT and ERK1/2, containing 21 measured lanes. Band rows were located manually and lane proposals reviewed. Common rectangles were supplied to both measurements. This convenience subset does not estimate success on an unselected image population.

Images were prepared as 8-bit inputs for application and reference decoding. The Python reference computed channel-mean intensity and the same local-background subtraction. We report relative differences for net signal and absolute differences for ratios to the first lane, avoiding a shared tolerance for quantities with different scales.

The 16-bit check used a generated membrane with six prescribed lanes and deterministic noise. Two TIFF outputs were produced using the same seed, dimensions and intensity design. Hash comparison confirmed identical files. They provide 12 measurements of six distinct constructed lanes, not two independent synthetic experiments. The fresh run called the fixed-version image decoder and measurement function. A previous rolling-ball component comparison is separate from this study’s local-background comparison and is not presented as a complete ImageJ end-to-end benchmark.

### Histology inputs and comparator audit

The functional corpus comprised 35 vendor teaching or product images: 12 H-DAB, four H&E, two trichrome and 17 multiplex or double-stain images processed through the fluorescence workflow. These were browser-sized example images, not calibrated whole slides. Successful execution was separated from quantitative agreement; independent references were not available for all 35 images or metrics.

The H-DAB area comparison covered eight images and ten ROIs: six whole-image regions and two pairs of subregions. The Python reference decoded pixels separately and performed stain separation and thresholding. During manuscript preparation, we found that it selected application ROIs by display name and silently substituted a whole-image result when a name was missing. Two counterstain ROIs had different labels in the configuration and application output. Both comparisons had consequently used the same whole-image percentage.

Further checks identified differences between Python and JavaScript rounding of a half-pixel ROI coordinate, and between continuous-background estimation in the original reference and the application’s clipping and histogram-based background rule. We retained the earlier records and corrected the comparator in stages. ROI matching now requires unique agreement of the recorded geometry and pixel count; an absent or ambiguous match raises an error. The independent reference was recalculated from the local source images. The fixed pathology source revision was also rerun on all 35 images. The final area comparison pairs fresh application and reference outputs, with explicit ROI geometry and pixel-count checks. Independent-threshold and fixed-application-threshold results were retained rather than selecting only the closer comparison.

### qPCR inputs and numerical checks

The main input was `RELATIVE_example.csv` from the Auto-qPCR example dataset [6], comprising four samples, five targets and three technical replicates per sample–target combination (60 wells). Three configurations exercised a two-group Welch path, an ANOVA/Dunnett path with two reference genes, and a Pfaffl path with prescribed efficiencies of 92–105%. A fourth used constructed Cq values for moderate effects. Configurations reuse inputs and are not independent biological datasets.

A Python reference recomputed summaries, expression values and probabilities. The comparator used an absolute tolerance of 10^−9^ for Cq-domain summaries and a relative tolerance of 10^−9^ for fold-change fields, scaling the difference by the larger absolute paired value with a 10^−12^ floor. Ordinary Welch and ANOVA probabilities used an absolute tolerance of 10^−8^. Printed Welch statistics were available only to two decimal places and used an absolute tolerance of 0.0051. Dunnett-adjusted probabilities used a relative tolerance of 0.02, with a 10^−300^ denominator floor, together with the bounds raw p ≤ adjusted p ≤ min(1, k × raw p), allowing 10^−12^ absolute slack; k is the number of treatment comparisons. The observed Dunnett errors were smaller than this tolerance (Figure 4). These criteria are not a claim that all outputs have nine or eight correct decimal places. Supplementary Material S1 preserves each field-level check and its comparison rule.

## Results

### Western blot signal agreement

In the fixed-version rerun, application and reference local-background net signals were equal at recorded precision for **all 21 lanes across four panels** (Figure 2). Figure 2 shows paired values against the identity line, a graphical first step in method comparison [27], and the absolute difference for each lane. Net signals and ratios were serialized as floating-point numbers without explicit decimal-place rounding; the zero difference was verified from these paired values rather than only the rounded error-summary fields. Ratios normalized to the first lane were also equal in the separate ratio table. Both sides used the same prepared TIFF inputs, whose hashes were recorded. Earlier comparison records had small nonzero discrepancies, but their prepared reference inputs could not be matched to the present input hashes; those records are retained as superseded audit evidence, not combined with this rerun. The result supports agreement for selected rectangles and prepared inputs, not all automatic proposals, acquisition formats or background modes.

**Figure 2.**
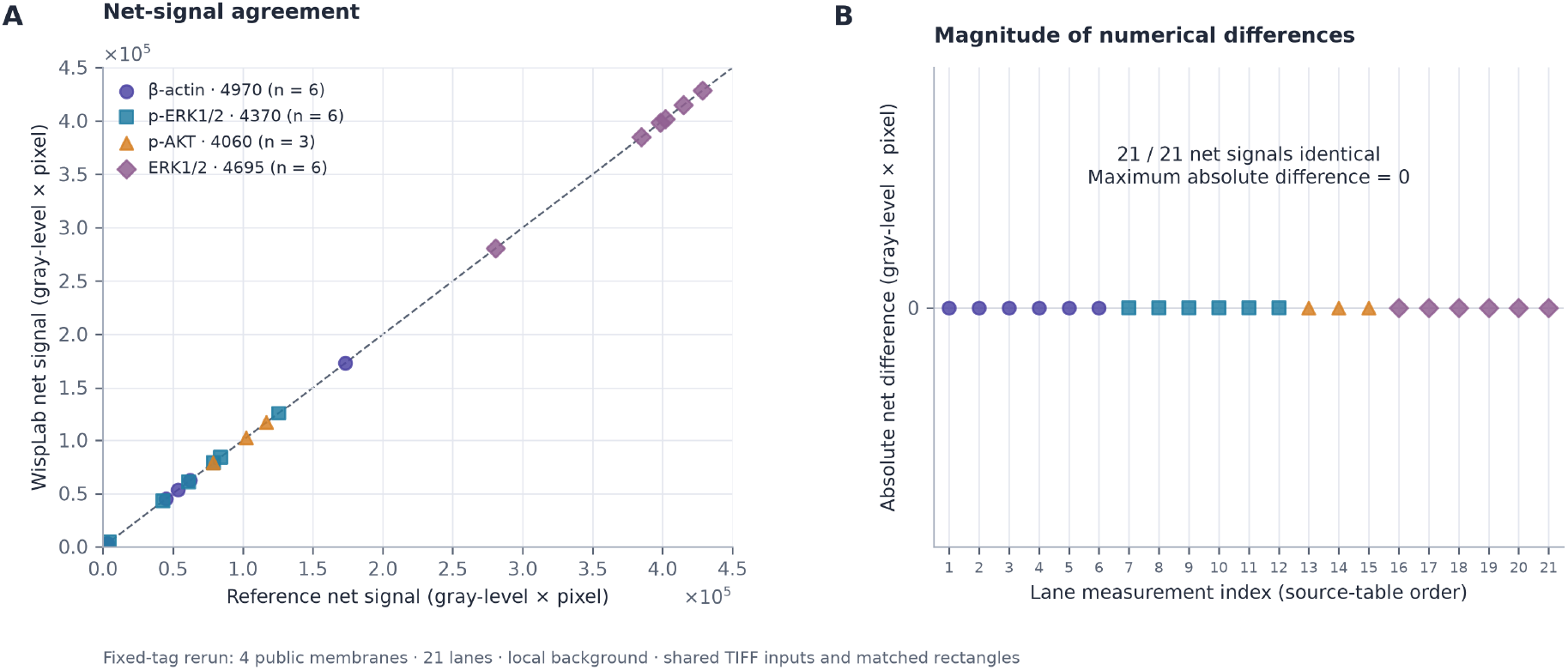
WB local-background net-signal agreement. A. Application and reference net signals for 21 lanes from four public panels; colors and shapes identify panels. Dashed line, identity. (B) Absolute differences for every lane, in source-table order. All paired net signals are equal in the serialized floating-point outputs, so all points lie at zero. Signal units are gray level multiplied by pixel count. Both implementations used the same prepared TIFF inputs and matched rectangles. These measurements are not independent biological replicates and do not benchmark automatic localization or other background modes. Data derive from the fixed-source rerun on 18 October 2026. Synthetic measurements are reported separately and are not pooled with these points.

In the fresh fixed-version 16-bit run, all 12 measurements matched their constructed reference values at recorded precision, with maximum absolute difference zero. Identical hashes confirmed that the two inputs represented one six-lane design. This exercises constructed decoding and measurement cases, not real 16-bit acquisition variability. Low-contrast loading-control panels and complex layouts remain reasons to inspect proposed regions.

### Histology comparison and correction of verification errors

The 35-image record supports execution of the exercised workflows; quantitative reference comparison is limited to H-DAB. The two previously mispaired application ROIs have positive fractions of 5.0019% and 8.9626%, rather than the duplicated whole-image value of 7.1151%. Their corresponding reference values change when coordinate rounding and preprocessing are aligned. Figure 3 presents the final comparator results; the staged corrections and earlier outputs are retained in the audit material.

**Figure 3.**
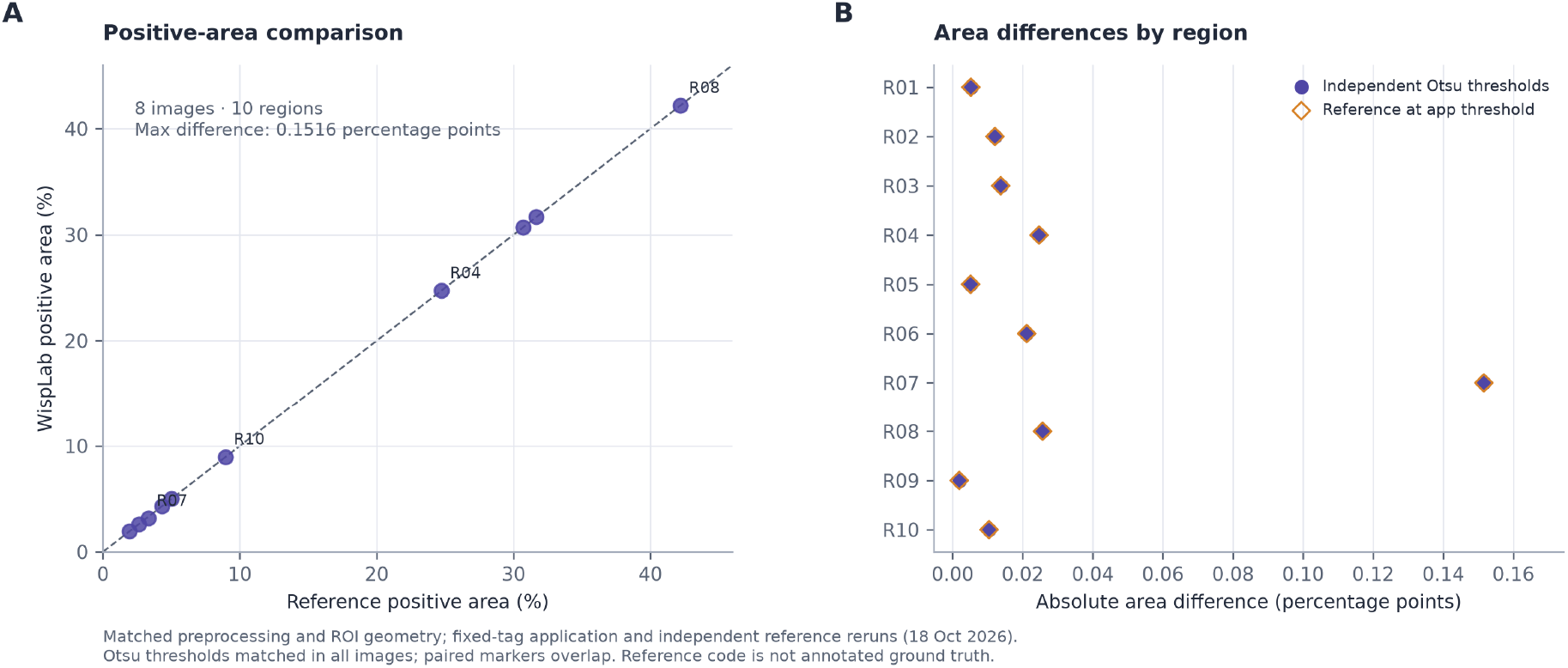
H-DAB positive-area agreement. A. Application and independent-reference positive-area percentages for ten regions from eight H-DAB images. Dashed line, identity. (B) Absolute differences using independently computed reference Otsu thresholds (filled circles) or application thresholds (open diamonds). After preprocessing and ROI alignment, thresholds were identical for all eight images, so paired markers overlap. The maximum difference was 0.151597992 percentage points at R07, the weak region of the step-5 image. All regions are retained. The figure uses fresh fixed-source application outputs and the corrected independent reference. It does not validate H-score, cell counts, H&E, trichrome or fluorescence measurements. Earlier comparator errors and intermediate records remain in Supplementary Material S1.

With aligned preprocessing and uniquely matched regions, the maximum absolute difference was **0.151597992 percentage points** across eight images and ten ROIs. All eight image-level Otsu thresholds matched, so the independent-threshold and fixed-application-threshold comparisons yielded the same area differences. The largest difference occurred in the weak region of the step-5 teaching image. The counterstain reference fractions were 5.0038% and 8.9522%, versus application fractions of 5.0019% and 8.9626%. These results concern agreement of matched calculations, not annotated biological truth. The superseded maximum of 2.1867 percentage points is not retained as a final result, and no isolated decoder cause is assigned to the remaining difference.

No independent accuracy claim is made for H&E, trichrome, nuclear counts, H-score or fluorescence colocalization. A historical threshold-calibration demonstration used a near-negative image as a pseudo-control. It was not a true unstained or no-primary control and is not used as accuracy evidence.

### qPCR agreement and deep-tail defect

A fresh run of source revision `4142b6078251d3973667bb59352a016ac0ebf3da` reproduced **422 passing checks across four configurations**, with no failed checks. This count includes repeated fields and qualitative assertions; it is not a count of independent experiments. Expression summaries and ordinary statistical probabilities passed their specified criteria (Figure 4). The Pfaffl configuration used prescribed efficiencies and does not validate efficiency estimation from amplification curves.

**Figure 4.**
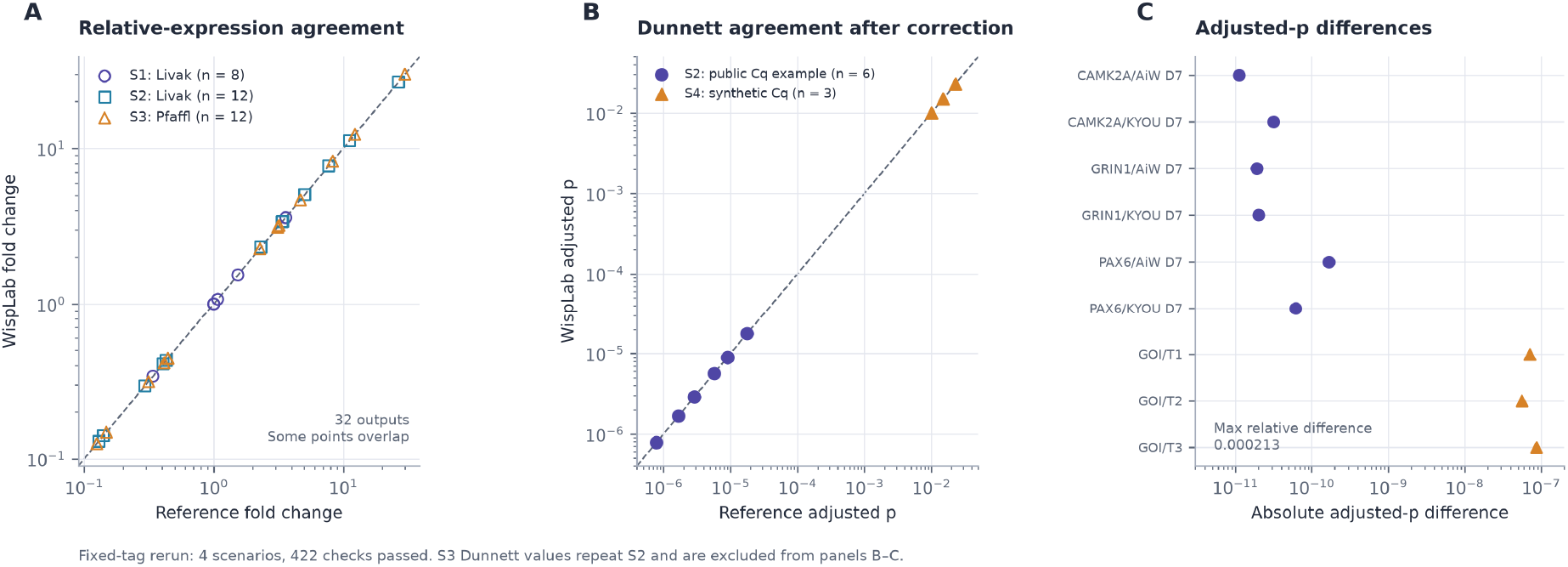
qPCR expression and adjusted-probability agreement. A. Thirty-two fold-change outputs from scenarios S1–S3 using the shared 60-well example with Livak or Pfaffl settings; some points overlap. Maximum absolute difference, 3.55 × 10^−15^. (B) Corrected Dunnett-adjusted probabilities for six contrasts from S2 and three from the synthetic S4 scenario. Dashed line, identity. (C) Absolute differences for those nine contrasts. Maximum relative difference, 2.125636791 × 10^−4^ (approximately 0.0213%); maximum absolute difference, 8.706212099 × 10^−8^. S3 repeats the S2 statistical contrasts and is omitted from B–C to avoid duplicating those values. The fixed-source rerun passed 422 checks across four scenarios (89, 133, 133 and 67); checks and outputs are not independent experimental observations. No pre-correction values are plotted. Technical wells provide computational examples, not new biological inference.

Original verification had exposed an application defect in strong-effect Dunnett comparisons. One documented adjusted probability, approximately 6.88 × 10^−13^, was below its corresponding unadjusted probability of approximately 1.61 × 10^−6^, contradicting the adjustment’s probability definition. The original quadrature missed a narrow tail contribution. The correction integrates the tail in the chi-square variable with adaptive truncation; the Python reference also required numerical refinement.

Post-fix comparisons and a regression case document the correction. For the illustrated strong-effect examples, the displayed significance category remained unchanged while the exported numerical probability changed substantially. This observation is limited to those examples and does not establish that every downstream conclusion from the earlier version was unaffected.

## Verification summary

**Table 1.**
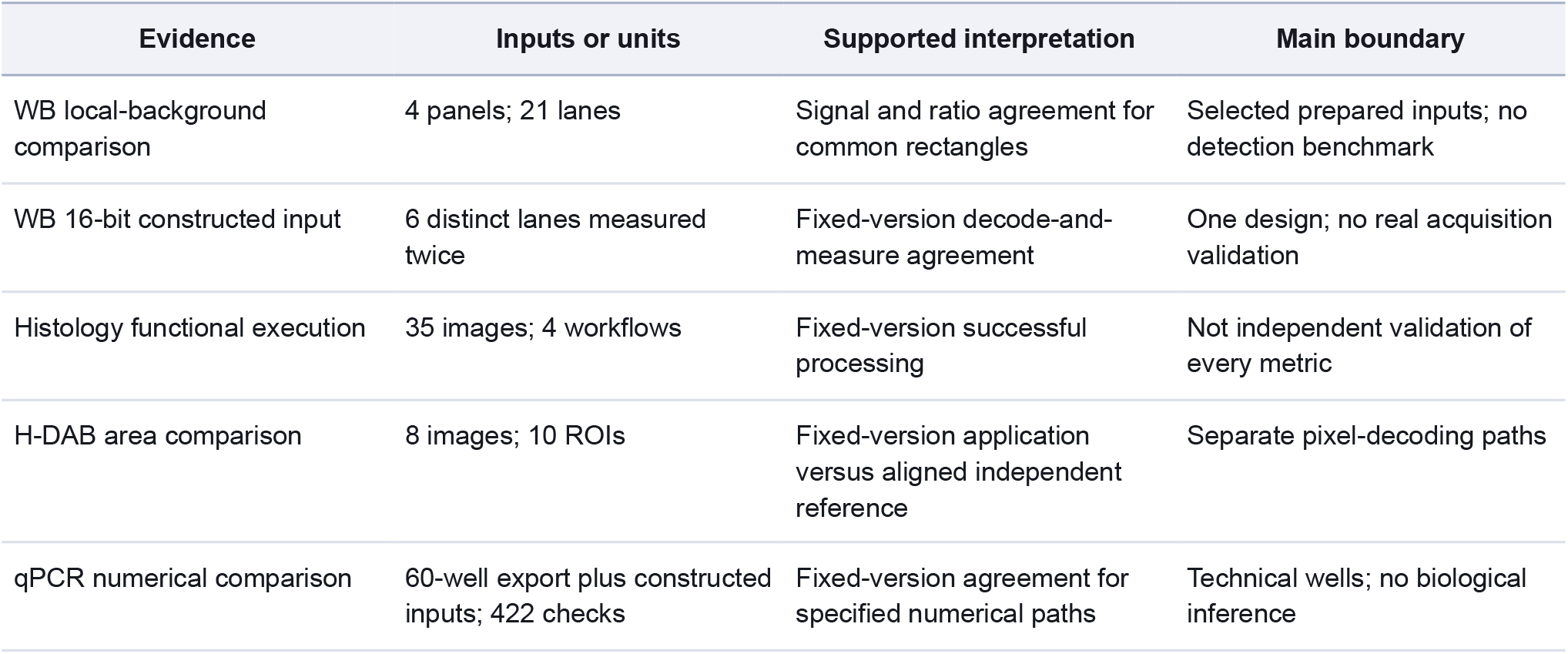
Verification denominators and boundaries. Inputs overlap between rows and should not be summed as independent validation experiments.

## Discussion and limitations

WispLab connects selected laboratory inputs to measurements and exported summaries. The contribution is an inspectable implementation with verification records, rather than a new quantification algorithm. Established tools remain appropriate alternatives and reference points [1,2,4-6]. Usability, speed, memory consumption and analytical superiority were not compared formally.

Related tools occupy complementary niches: CellProfiler addresses high-throughput image-analysis pipelines [3], IHC Profiler automates ImageJ-based IHC scoring [5], blotRig targets Western blot experimental design and mixed-model analysis [12], and Click-qPCR provides guided interactive processing of qPCR tables [28]. WispLab differs in linking Western blot, staining-image and qPCR workflows in bilingual browser tools with exported parameters and reference-comparison records. A systematic feature comparison was not performed, and no analytical superiority is claimed.

Failures arose in both sides of the comparison. A probability-bound violation motivated a qPCR application correction. Incorrect ROI matching and inconsistent preprocessing required correction of the histology reference. These cases show why provenance, input alignment and checks of the comparator are needed alongside a reported tolerance. Matching after independently chosen thresholds alone is insufficient to establish equivalence of the underlying stain channel.

The selected image examples do not establish robustness across scanners, acquisition settings, staining protocols or laboratories. Area agreement does not validate nuclear segmentation or clinical scoring. Original multichannel fluorescence inputs and appropriate references are needed before claiming colocalization accuracy. A calibrated image cohort, expert annotations and QuPath-based comparisons would strengthen future validation.

Western blot region selection was supplied to the comparison, and local background was the evaluated correction. Further work should include original 16-bit acquisition files, varied layouts and full end-to-end comparisons with established tools. Rolling-ball component tests must remain distinct from those studies. The duplicated synthetic files cannot be counted as independent coverage.

For qPCR, agreement concerns computations applied to selected example wells. Experimental validity still depends on specificity, efficiency estimation, controls, biological replicate structure and reporting [20,21]. Technical triplicates do not by themselves justify population-level inference. Independent comparison did not cover all parser layouts or quality-control heuristics. Dunnett quadrature was evaluated over selected examples, not proven uniformly accurate for all degrees of freedom and effect sizes.

Some source images cannot be redistributed. URLs and acquisition records improve traceability but cannot guarantee future availability. Generated inputs and redistributable numerical examples provide a reproducible subset; the image comparisons retain an external-input dependency. The reproducibility report separates fresh fixed-source reruns from historical results with incomplete provenance. The reruns reused installed dependencies, so clean installation from the archived dependency locks remains a separate check.

## Supporting information

Supplementary Material S1: verification records, scripts and figure source data

## Availability

All source code for the research versions is openly available from the repositories below, with licence notices in the repositories. Third-party components and input materials retain their own terms.

| Component | Repository | Version DOI | Paper-tag commit (prefix) |
| --- | --- | --- | --- |
| WB and portal | blot-workbench | 10.5281/zenodo.23439309 | 6bc90cec4422 |
| IHC / staining / IF | pathology-workbench | 10.5281/zenodo.23439313 | dd41bae4bc27 |
| qPCR | pcr-workbench | 10.5281/zenodo.23439315 | 4142b6078251 |

All three releases use `wisplab-paper-2026-10-17`. Applications are linked from wisplab.top, with endpoints at wb.wisplab.top, pathology.wisplab.top and pcr.wisplab.top. Live deployments may be newer than the paper releases.

**Supplementary Material S1** supplies the verification scripts, comparison records, corrected comparator, plotting code, numerical figure-source tables and reproducibility instructions. It accompanies the manuscript as a separate archive; the three code-version DOIs above identify the archived application sources. Vendor images and overlays are excluded from redistribution. The fixed PCR tag incorrectly labels the Auto-qPCR examples as MIT; S1 preserves their upstream GPLv3 licence and provenance. The Auto-qPCR example export and RDML parser fixtures are separately identified by provenance; test-material permissions are not generalized to application code.

## Author contributions

Yuan Gao and Sumin Sun jointly conceived and designed the work, directed the software development and verification programme, curated the verification inputs, interpreted the comparison records, and wrote and revised the manuscript. Both authors read and approved the final version and agree to be accountable for all aspects of the work.

## Data provenance and ethics

The verification used publicly accessible example data, vendor teaching or product images, and generated test inputs. No new participant recruitment or animal experiments were performed for the verification reported here. Public accessibility does not imply permission to redistribute third-party images.

## AI assistance

OpenAI Codex assisted with software development, manuscript drafting and editing, reference checking, and preparation of analysis and plotting scripts. Suggestions and numerical claims were checked against source files, comparison records and primary publications during preparation. The authors are responsible for the final manuscript and its conclusions.

## Abbreviations

ANOVA: analysis of variance
Cq: quantification cycle
H-DAB: haematoxylin–diaminobenzidine
H&E: haematoxylin and eosin
IF: immunofluorescence
IHC: immunohistochemistry
MIQE: Minimum Information for Publication of Quantitative Real-Time PCR Experiments
NSFC: National Natural Science Foundation of China
OD: optical density
ONNX: Open Neural Network Exchange
qPCR: quantitative real-time polymerase chain reaction
ROI: region of interest
TIFF: tagged image file format
WB: Western blot

