## Supplementary Material S1: verification records, scripts and figure source data for "WispLab: open-source browser workbenches for laboratory image and qPCR analysis with reference-based verification": FIGURE-NOTES.md

# WispLab preprint figures

Prepared 18 October 2026. These are data-driven manuscript figures, not additional biological experiments. The figures contain no vendor source images, overlays or screenshots. Figure 1 is an original vector workflow diagram. Figures 2–4 plot numerical records. All labels are in English, and each figure is supplied as PDF, editable-text SVG and 300-dpi PNG.

## Files and reproducibility

| Figure | Output basename | Numerical source table |
| --- | --- | --- |
| 1. Workflow and verification scope | `figure1_workflow` | Conceptual workflow; no measured values beyond recorded coverage |
| 2. WB net-signal agreement | `figure2_wb_agreement` | `figure2_wb_source.csv` |
| 3. IHC positive-area agreement | `figure3_ihc_area` | `figure3_ihc_source.csv` |
| 4. qPCR agreement | `figure4_pcr_agreement` | `figure4_pcr_fold_change_source.csv`, `figure4_pcr_dunnett_source.csv` |

`figure-data-summary.json` contains computed counts, error summaries, input SHA-256 values and plotting package versions. `source-data/source-manifest.json` maps preserved input snapshots to their original paths and distinguishes superseded historical records from fixed-tag application/reference reruns and the IHC audit corrections. `pcr_check_counts.csv` gives the four recorded scenario totals. `wb_synthetic_16bit_source.csv` preserves the synthetic WB records without treating duplicate files as independent evidence.

From the manuscript directory, use an isolated Python environment:

```sh
python3 -m venv .venv-figures
.venv-figures/bin/python -m pip install -r figures/requirements-plotting.txt
.venv-figures/bin/python scripts/plot_figures.py
```

The script locates the source snapshots relative to itself and can run from any working directory. Optional arguments `--source-dir` and `--output-dir` accept alternative directories. Plot regeneration does not rerun the application engines or independent numerical references. The figures use Matplotlib 3.11.2 and NumPy 2.5.4. No smoothing, down-sampling or omission based on error magnitude is performed. Scatter plots retain overlapping observations. No confidence intervals or biological error bars are appropriate for these deterministic paired calculations.

## Suggested figure legends

### Figure 1. WispLab workflow and scope of the recorded verification

The three browser workbenches accept Western-blot images, microscopy images or qPCR Cq tables. Users inspect analysis settings and regions and export results. The default assay computations run in the browser; optional external AI services fall outside this default local flow and may transmit user-selected data. The lower panel summarizes the numerical comparisons used here: four public WB membranes comprising 21 measured lanes, eight DAB/IHC images comprising ten analysis regions, and a 60-well public qPCR example supplemented by a synthetic numerical scenario. These are implementation checks with independent Python reference code, not a clinical validation or a comparison of biological accuracy.

### Figure 2. Agreement of local-background WB net signals from a fixed-version rerun

(A) Application and independent-reference net signals for 21 measured lanes from four public membrane images. Colors and marker shapes identify the source membranes; sample sizes denote lanes, not independent biological replicates. Dashed line, identity. (B) Absolute application–reference differences for all 21 lane measurements, shown in source-table order. All paired net signals are exactly equal in the recorded full-precision outputs; every marker is at zero, and only the zero y-axis tick is shown. Net signal units are gray level multiplied by pixel count. Both independently implemented calculations used the same freshly converted TIFF inputs and matched measurement rectangles, with local background subtraction after target-band-row selection. Exact equality here is restricted to these inputs and rectangles. This figure does not test rolling-ball subtraction, fully automatic lane localization, biological ground truth, or equivalence across arbitrary image decoders and acquisition systems.

Source: `manuscript/reproducibility/wb-real-pinned-20261018/reference_comparison.json`, retained as `source-data/wb_reference_comparison_pinned_20261018.json`; rerun dated 18 October 2026, using application commit `6bc90cec4422d3e414629ce3d78b639c75279588`. Only its 21 real-image lanes are used in this figure. The source contains complete-precision net signals. Historical inputs had an unresolved staging/provenance mismatch; the old 3.17125 × 10⁻⁴ relative difference and 54.59 absolute difference are therefore **not** current results. The historical JSON remains in `source-data/wb_reference_comparison.json` for audit only.

The separate synthetic source is `manuscript/reproducibility/fresh-20261018/wb/comparison_report.json`, retained as `source-data/wb_synthetic_comparison_pinned_20261018.json`. Its full-precision application/truth values, rather than the rounded display fields embedded in the real-image comparison JSON, generate `wb_synthetic_16bit_source.csv`. This record has twelve measurements across two byte-identical six-lane files: **six unique constructed lanes**, not two independent synthetic images. All twelve recorded absolute differences are zero. Synthetic observations are not pooled with the 21 public-image points. Existing installed dependencies were reused for the reruns; this is not a clean dependency-installation test.

### Figure 3. Positive-area agreement with an independent reference after preprocessing and ROI alignment

(A) Application and independent-reference positive-area percentages for ten regions from eight DAB/IHC images. Dashed line, identity. (B) Absolute area differences in percentage points for every region, evaluated using the reference's independently computed Otsu threshold (filled circles) and the application's threshold (open diamonds). After matching preprocessing conventions, Otsu thresholds were identical for all eight images; the paired markers therefore overlap. Both comparisons have a maximum absolute difference of 0.151597992 percentage points, at R07 (the weak region of `leica_step5_full.jpg`). All points are retained. This comparison is restricted to positive area in DAB/IHC images and does not validate H-score, cell counts, H&E, trichrome, fluorescence or colocalization metrics.

Source used in the figure: `manuscript/reproducibility/record-audit-pinned-20261018/ihc-corrected-records.json`, retained as `source-data/ihc_reference_area_comparison_pinned_20261018.json`. On 18 October 2026 the independent Python reference was rerun on eight source images. Its conventions were aligned to the application for optical-density endpoint clipping, truncation of negative stain concentrations, Float32 intermediate channels, 256-bin integer-percentile background estimation, and JavaScript-style ROI rounding. ROI names and geometry were explicitly matched against historical application configuration; the reference did not import the application implementation. This establishes a comparison of separately implemented matched calculations, not validation against annotated biological truth. The application was subsequently rerun from the fixed research commit `dd41bae4bc27fda0b6bf1135b802e4af41ae7316`; the final plot uses that rerun, with runtime provenance recorded. All 35 generated image-analysis JSON outputs matched the historical numerical outputs; the quantitative comparison remains the eight-image, ten-region DAB/IHC subset. Existing installed dependencies were reused, so this was not a clean dependency-installation test. The earlier historical-output-only comparisons remain available for audit.

The original comparison incorrectly fell back to a whole-image percentage (7.1151%) for both regions of `leica_step13_full.jpg` when reference and application ROI names differed. The audit corrected that fallback and a Python/JavaScript half-pixel rounding difference, then aligned the preprocessing conventions above. The final paired values are reference 5.003779289% versus application 5.001889645% for R09 and reference 8.952191988% versus application 8.962585034% for R10. The initial maxima of 2.1867/5.3708 percentage points and intermediate geometry-only maxima of 0.550587972/4.544321696 percentage points describe superseded comparisons and must not be presented as final matched-method validation outcomes. Both historical and intermediate JSON snapshots remain in `source-data/` for the audit trail, with `used_in_figures: false` in the manifest. No image-decoder cause is inferred from the remaining discrepancy.

### Figure 4. qPCR expression and Dunnett-adjusted-p agreement with an independent numerical reference

(A) Thirty-two fold-change outputs across scenarios S1–S3, calculated from a shared 60-well public example using Livak or efficiency-corrected Pfaffl settings. Counts are reported outputs, not independent experimental replicates, and some points overlap. The maximum absolute fold-change difference was 3.55 × 10⁻¹⁵. (B) Dunnett-adjusted p values after the numerical correction, compared with the independently implemented quadrature reference. Six contrasts from the public Cq example (S2) and three from the synthetic Cq scenario (S4) are shown. Dashed line, identity. (C) Absolute differences for the same nine contrasts. The maximum relative adjusted-p difference was 2.125636791 × 10⁻⁴ (approximately 0.0213%); the maximum absolute difference was 8.706212099 × 10⁻⁸. S3 repeats the statistical contrasts from S2 and is excluded from panels B–C to avoid plotting the same p values twice. The fixed-tag rerun report contains 422 passing checks across four scenarios; these are implementation checks, not 422 independent observations. No pre-correction values are plotted because the selected source report contains the corrected results.

Source: `manuscript/reproducibility/fresh-20261018/pcr/comparison_report.json`, retained as `source-data/pcr_comparison_report_pinned_20261018.json`. This application/reference rerun on 18 October 2026 used the fixed research commit `4142b6078251d3973667bb59352a016ac0ebf3da` with existing installed dependencies. The historical report remains in `source-data/pcr_comparison_report.json` for audit only. This is not a clean dependency-installation test. The fold-change panel uses checks whose field ends in `.foldChange`; Dunnett panels use `.dunnettP`, `ref_quad`, and the recorded relative difference. The Dunnett CSV includes all 15 recorded S2/S3/S4 checks with a `plotted` field so that the six excluded repeated S3 values remain traceable. The report's 422 checks comprise 89 in S1, 133 in S2, 133 in S3 and 67 in S4. These plots do not establish the biological validity of the example's sample labels or demonstrate general accuracy for untested statistical designs.

## Validation and limitations of this figure package

- The plotting script asserts 21 public WB lanes, 12 recorded synthetic lane measurements (six unique constructed lanes), ten IHC regions/eight images, 32 fold-change outputs, nine plotted Dunnett contrasts and zero failures in the selected fixed-tag qPCR rerun report.
- Original numerical snapshots and the fixed-tag rerun snapshots have SHA-256 provenance. The obsolete and intermediate IHC snapshots are excluded from plotted data and explicitly retained for audit only.
- These plots are implementation agreement displays. Small paired differences do not establish biological truth, superiority, usability, reproducibility on every operating system, or equivalence to ImageJ/QuPath end-to-end workflows.
- The graph of equal-threshold IHC discrepancies is deliberately retained. The independently computed thresholds happen to match in this tested subset; this does not establish universal mask equivalence or biological truth.
- No new biological images or annotations were generated; application/reference reruns are recorded explicitly. The conceptual workflow diagram makes no claim that every available product feature has been independently benchmarked.
