## Supplementary Material S1: verification records, scripts and figure source data for "WispLab: open-source browser workbenches for laboratory image and qPCR analysis with reference-based verification": README.md

# Supplementary Material S1: WispLab numerical verification

Prepared 18 October 2026 for the accompanying manuscript. This local archive has
not yet been uploaded. The three application source versions are fixed at tag
wisplab-paper-2026-10-17:

- WB: 6bc90cec4422d3e414629ce3d78b639c75279588, doi:10.5281/zenodo.23439309
- Pathology: dd41bae4bc27fda0b6bf1135b802e4af41ae7316, doi:10.5281/zenodo.23439313
- PCR: 4142b6078251d3973667bb59352a016ac0ebf3da, doi:10.5281/zenodo.23439315

## Contents and reproduction

reproducibility/REPRODUCIBILITY.md provides exact commands, dependency limitations,
source/input hashes and the correction audit. claim-evidence.json identifies the
current records. Historical/intermediate comparisons are retained as audit only.
The fresh runs reused installed dependencies; no clean lockfile installation or
independent-machine reproduction is claimed. Acquisition manifests/configurations
identify external inputs. Vendor images, overlays and synthetic TIFFs are excluded.
Image reruns require lawful access to the specified external images; source URLs
may change. Synthetic TIFFs can be regenerated.

To regenerate the four figures from included numerical snapshots, from this root:

    python3 -m venv .venv
    .venv/bin/python -m pip install -r figures/requirements-plotting.txt
    .venv/bin/python scripts/plot_figures.py

This plots saved records and does not rerun application engines. See
figures/FIGURE-NOTES.md for scope and legends. All plotted points are retained;
repeated S3 Dunnett contrasts are identified rather than counted twice.

## Provenance and third-party terms

Read reproducibility/third-party-notices/ before reusing source-derived records.
The upstream Auto-qPCR example licence is distinct from WispLab application-code
licences. This package makes no blanket licence claim over third-party materials.
No upstream RDML fixture or vendor image is bundled. Application sources remain
at the fixed repositories/archives above and are not copied into this ZIP.

Public-copy and package manifests record SHA-256 checksums. Local filesystem home
and temporary prefixes were redacted in designated copies; original evidence is
retained by the preparer. Hashes for redacted copies can differ from original-run
hashes. This transformation changes paths, not numerical scientific values.
