## Supplementary Material S1: verification records, scripts and figure source data for "WispLab: open-source browser workbenches for laboratory image and qPCR analysis with reference-based verification": README.md

# Image-source manifests and configuration

This directory contains source URLs, local-input checksums and scope descriptions,
not vendor images. Public access to a URL is not a redistribution license.
The publisher's terms apply, and links or image renditions can change.

## Western blots

`wb-MANIFEST.json` preserves the ordered ten-entry list consumed by the historical
harness. Four successful real-image panels are explicitly flagged as the
21-lane comparison subset; other entries document attempted candidates and the
historical synthetic filenames, not additional successful comparisons. To rerun
the unchanged harness, arrange the lawfully obtained input files at the manifest's
relative paths and copy this manifest to `WB_CORPUS/MANIFEST.json`.

Panel-selection coordinates and candidate lane counts are already supplied in
`scripts/wb_real_harness.ts`. Final band/background rectangles are in
`wb-real-pinned-20261018/app_results.json`; all source images remain external.
The independent synthetic generator does not require vendor images and is
available separately in `scripts/wb_synthetic.ts`.

## IHC, staining and IF

`ihc-35-input-sources.json` lists exactly the 35 fresh-run inputs, their relative
paths, SHA256, source pages and observed direct-image URLs. The 12 DAB direct URLs
come from the original source manifest. For the remaining images, responsive
image URLs were extracted from saved vendor HTML; these are observed candidates,
not a claim that every URL resolves to the exact locally tested byte sequence.
Use the checksums to establish identity. No image was downloaded during this
manifest preparation.

The eight-image quantitative subset is specified by the `CONFIG` literal in
`scripts/ihc_reference_aligned.py`. Scene presets, full-image/subregion selection
and fractional ROI coordinates are in `scripts/ihc_harness.mjs`. The final
reference report records resolved pixel rectangles and uniquely matched pixel
counts. This makes the comparison configuration inspectable without duplicating
the vendor image material.
