## Supplementary Material S1: verification records, scripts and figure source data for "WispLab: open-source browser workbenches for laboratory image and qPCR analysis with reference-based verification": REPRODUCIBILITY.md

# WispLab preprint: pinned-source reproduction, 18 October 2026

This directory contains actual reruns, not just recalculated errors from old
application outputs. No production repository, historical validation result,
license, or previously prepared public deposit was changed.

## Current evidence to use in the manuscript

| Module | Pinned source commit | Fresh run and scope | Result |
|---|---|---|---|
| qPCR | `4142b6078251d3973667bb59352a016ac0ebf3da` | CSV parsing + application engine + independent Python reference; 4 scenarios | 422 checks passed, 0 failed. Among 15 adjusted p comparisons, maximum relative difference 0.00021256367909340264. |
| WB real images | `6bc90cec4422d3e414629ce3d78b639c75279588` | Same newly converted TIFF input for both implementations; manually selected panels + proposed lanes + local-background measurement; 4 images/21 lanes | All net signals and all first-lane-normalized ratios exactly equal in this run. |
| WB synthetic 16-bit | same WB commit | Generate, decode and measure a constructed TIFF twice | 12 measurements, **one unique image and six unique lanes**; all net signals exactly equal to constructed truth. |
| IHC/staining/IF execution | `dd41bae4bc27fda0b6bf1135b802e4af41ae7316` | Isolated Vite + Chromium runs of official engines, 35 images | 35 completed, 0 failed: 12 DAB, 17 IF, 4 H&E, 2 trichrome. This is functional coverage, not independent metric validation. |
| DAB positive area | same pathology commit | Fresh browser outputs + fresh independently written Python reference, aligned preprocessing and ROI definitions; 8 images/10 ROI | Maximum absolute difference **0.1515979919479098 percentage points**. Otsu thresholds match on all eight images; both independent-threshold and same-threshold comparisons therefore have this maximum. |

`claim-evidence.json` and `claim-evidence.tsv` map these claims to files.
Use `record-audit-pinned-20261018/ihc-corrected-records.json` for IHC figures,
`wb-real-pinned-20261018/reference_comparison.json` **8-bit rows only** for WB,
`wb-real-pinned-20261018/ratio_comparison.csv` for normalization ratios, and
`fresh-20261018/pcr/comparison_report.json` for qPCR.
The canonical synthetic result is `fresh-20261018/wb/comparison_report.json`;
the synthetic rows in the real-image reference report retain historical
two-decimal display formatting for truth values and should not be used as
the synthetic precision table.

## What was and was not reproduced

The source trees were extracted with `git archive` at the full commits above
into disposable directories; production code was not modified. Existing local
`node_modules` and the installed Playwright browser were reused. This is a
**source-isolated rerun, not a fresh dependency-installation or independent
machine reproduction**. Reports preserve source-file hashes, harness hashes,
actual package versions where collected, input hashes and timestamps.

The runtime was macOS 27.0.1 arm64, Node v22.22.0 and Python 3.9.6. qPCR used
tsx 4.23.15/TypeScript 6.0.3; WB used tsx 4.23.15, tiff 7.1.3, fast-png 8.0.0
and fflate 0.8.3. IHC used the versions recorded in its engine run report.

The Python reference code does not import production computation functions.
Matching the computation specification is necessary for a numerical consistency
comparison; it does not establish that the specification is biologically valid.
IHC reference input decoding uses macOS `sips`, while application decoding uses
Chromium. Decoded pixel arrays were not forced identical. Residual differences
were not experimentally attributed to one decoder or floating-point source.

No new biological experiment, clinical validation, true negative-control
validation, independent nuclei/H-score/IF-colocalization metric benchmark,
ImageJ end-to-end comparison or QuPath comparison was performed. WB exact
agreement applies to the tested pixels, rectangles, arithmetic and local
background definition; it is not a universal error bound.

## Historical-record corrections and provenance

1. Historical IHC `compare_reference.py` looked up ROI display names and silently
   substituted `overall` when a name was absent. For step13, the reference labels
   were `A(深复染)`/`B(浅复染)` whereas application outputs used `A(弱)`/`B(强)`.
   Both sides were incorrectly compared with the same whole-image 7.1151% value.
2. Python `round(10.5)` yielded 10, while browser `Math.round(10.5)` yielded 11.
   The step13 reference rectangle was shifted by one pixel vertically. New
   references use `floor(x + 0.5)` for these nonnegative coordinates, with strict
   geometry and pixel-count matching; missing or ambiguous matches raise errors.
3. The old reference applied a continuous 5% percentile to concentrations before
   negative-value clipping. The application clips negative concentrations first
   and uses a nearest-integer 256-bin histogram percentile. The aligned reference
   additionally matches channel-OD endpoint clipping and Float32 storage.
4. The retained audit sequence is: original 2.1867 pp maximum / 5.3708 pp at the
   application threshold; name + geometry correction 0.550587972 pp / 4.544321696 pp;
   preprocessing alignment 0.151597992 pp / 0.151597992 pp. The first two stages
   are audit history, not the final accuracy claim. All stages are retained.
5. The new pinned IHC run reproduced all 35 historical numerical JSON outputs
   exactly. Historical output files themselves still have no embedded runtime
   commit and their old provenance is **unknown**, despite README declarations.
   New source provenance comes from the new run, not from retroactively assigning
   a commit to old files.
6. The new pinned WB run reproduced every historical application net value and
   rectangle exactly. Its independent reference now reads the same newly
   prepared TIFFs and agrees exactly. Historical stored `wb/staged` TIFF file
   hashes differ from the new prepared inputs; their preparation is not fully
   traceable. A file-hash difference alone does not establish a pixel difference
   or prove the cause of the earlier nonzero reference discrepancy. Retain the
   historical 0.000317125 result only as historical audit evidence.
7. The old synthetic generator resets the same random seed for both file names.
   Fresh file hashes confirm that the two TIFFs are byte-identical. Count six
   unique constructed lanes, repeated twice, not 12 independent lanes.

## Reproduction commands

Run these from this directory. Replace example paths with local paths; each output
directory must be new. The required repositories must contain the pinned Git
objects, and installed Node dependencies must be available. No network download
or publishing action is performed by these scripts.

```sh
python3 scripts/run_reproduction.py \
  --pcr-repo /path/to/pcr-workbench \
  --wb-repo /path/to/blot-workbench \
  --output /path/to/new-pcr-and-synthetic-run

python3 scripts/run_wb_real_pinned.py \
  --repo /path/to/blot-workbench \
  --corpus /path/to/WB-corpus \
  --output /path/to/new-wb-real-run

python3 scripts/run_ihc_pinned.py \
  --repo /path/to/pathology-workbench \
  --ihc-root /path/to/IHC-corpus \
  --output /path/to/new-ihc-engine-run

python3 scripts/ihc_reference_aligned.py \
  --ihc-root /path/to/IHC-corpus \
  --app-results /path/to/new-ihc-engine-run/results_m3 \
  --output /path/to/new-ihc-reference-run

python3 scripts/audit_records.py \
  --validation-root /path/to/original-validation-records \
  --ihc-root /path/to/IHC-corpus \
  --app-results /path/to/new-ihc-engine-run/results_m3 \
  --app-run-metadata /path/to/new-ihc-engine-run/run-report.json \
  --ihc-reference-report /path/to/new-ihc-reference-run/reference_area_comparison.json \
  --output /path/to/new-record-audit
```

The optional `--pcr-node-modules` and `--wb-node-modules` flags on
`run_reproduction.py`, and `--node-modules` on `run_ihc_pinned.py`, allow dependency
directories installed elsewhere. The PCR fixture is read from the pinned
repository, not a mutable working-tree file. WB and IHC vendor images are local
inputs and are **not included** in this archive. Corpus manifests/source URLs and
lawful access to the specified inputs are still required for image reproduction.
The numeric record audit can be inspected without redistributing source images.
The included `datasets/` directory supplies the exact selected source pages,
observed image URLs, checksums and reconstruction instructions; it is not an
image redistribution grant.

The IHC audit intentionally fails on the uncorrected historical geometry if
`--ihc-reference-report` is omitted. This prevents a silent return to the original
comparison error. `ihc_reference.py` retains the intermediate geometry-only
reference; use `ihc_reference_aligned.py` for the final result.

## Archive preparation

Use `archive-allowlist.json`, not a recursive copy of the whole directory.
It includes scripts, numerical JSON/CSV/TSV, documentation and logs. It excludes
all image formats, `node_modules`, Python caches and temporary directories.
The synthetic image can be reconstructed with the generator and need not be
distributed as TIFF. No original vendor image or derived overlay is present in
the allowed payload.

```sh
python3 scripts/prepare_public_copy.py --output /path/to/new-public-reproduction-copy
```

That command creates a new local copy only. It replaces the local user's home
prefix in text with `<LOCAL_HOME>` and records file hashes for the public copy.
It neither changes original evidence nor publishes to Zenodo/bioRxiv. Public
copy hashes differ after this documented path de-identification. The existing
2026-10-17 deposit ZIP remains unchanged. Copyright/license review of source
code, upstream fixtures and source-image acquisition remains separate.

The Auto-qPCR fixture source and complete upstream license have now been verified
at fixed revision `d881ebd01667766882f90de850bfc26d93c54e27`. That upstream uses
GPL version 3, not the MIT attribution in older local notes. See
`third-party-notices/README.md`, the unchanged `Auto-qPCR-LICENCE.txt`, and the
byte-identity/provenance JSON. This data-material notice does not relicense the
WispLab application repositories.
