## Supplementary Material S1: verification records, scripts and figure source data for "WispLab: open-source browser workbenches for laboratory image and qPCR analysis with reference-based verification": audit_records.py

```
#!/usr/bin/env python3
"""Audit historical records. DOES NOT rerun image analysis or prove engine SHA.

IHC ROI correspondence is reconstructed from both archived harnesses' geometry,
then checked against output pixel counts. Missing/ambiguous matches fail closed.
"""
import argparse
import ast
import hashlib
import json
import math
from pathlib import Path
import re

def sha(p):
    return hashlib.sha256(p.read_bytes()).hexdigest()

def load(p):
    return json.loads(p.read_text())

def main():
    ap=argparse.ArgumentParser()
    ap.add_argument('--validation-root',type=Path,required=True)
    ap.add_argument('--ihc-root',type=Path,required=True)
    ap.add_argument('--ihc-reference-report',type=Path,help='Optional fresh, corrected reference output. Without it the historical geometry mismatch intentionally fails.')
    ap.add_argument('--app-results',type=Path,help='Optional fresh pinned IHC results_m3 directory')
    ap.add_argument('--app-run-metadata',type=Path,help='Pinned application engine run-report.json')
    ap.add_argument('--output',type=Path,required=True)
    a=ap.parse_args()
    if a.output.exists():
        raise SystemExit('Output directory must not already exist.')
    a.output.mkdir(parents=True)
    pcr_path=a.validation_root/'pcr/comparison_report.json'
    wb_path=a.validation_root/'wb/reference_comparison.json'
    ihc_path=a.ihc_root/'comparison/reference_area_comparison.json'
    ref_script=a.ihc_root/'compare_reference.py'
    app_script=a.ihc_root/'run_m3.mjs'
    # Read CONFIG as a Python literal, never execute historical code.
    tree=ast.parse(ref_script.read_text())
    configs=[ast.literal_eval(n.value) for n in tree.body if isinstance(n,ast.Assign) and any(isinstance(t,ast.Name) and t.id=='CONFIG' for t in n.targets)]
    assert len(configs)==1,'Missing/ambiguous reference ROI configuration'
    config=configs[0]
    text=app_script.read_text()
    expected_selector="f.startsWith('leica_step') && !['leica_step14_full.jpg', 'leica_step15_full.jpg'].includes(f)"
    assert expected_selector in text, 'Application ROI selector changed; inspect manually'
    patt=r"\{\s*fx:\s*([\d.]+),\s*fy:\s*([\d.]+),\s*fw:\s*([\d.]+),\s*fh:\s*([\d.]+),\s*name:\s*'([^']+)'\s*\}"
    app_geom={}
    for fx,fy,fw,fh,name in re.findall(patt,text):
        assert name not in app_geom,'Ambiguous application geometry'
        app_geom[name]=dict(zip(['fx','fy','fw','fh'],map(float,[fx,fy,fw,fh])))
    assert set(app_geom)=={'整图','A(弱)','B(强)'},'Unexpected application ROI literals'
    inputs={str(p):sha(p) for p in [pcr_path,wb_path,ihc_path,ref_script,app_script]}
    corrected=[]
    reference_path=a.ihc_reference_report or ihc_path
    inputs[str(reference_path)]=sha(reference_path)
    historical={(r['file'],r['roi']):r for r in load(ihc_path)['rows']}
    for old in load(reference_path)['rows']:
        path=(a.app_results or a.ihc_root/'results_m3')/'dab'/old['file'].replace('.jpg','.json')
        inputs[str(path)]=sha(path)
        data=load(path)
        refs=[r for r in config[old['file']] if r['name']==old['roi']]
        assert len(refs)==1,'Missing/ambiguous reference ROI'
        geom={k:refs[0][k] for k in ['fx','fy','fw','fh']}
        # Python reference uses round; the browser harness uses Math.round.
        pyrect=[round(geom[k]*data[axis]) for k,axis in [('fx','width'),('fy','height'),('fw','width'),('fh','height')]]
        jsrect=[math.floor(geom[k]*data[axis]+0.5) for k,axis in [('fx','width'),('fy','height'),('fw','width'),('fh','height')]]
        if a.ihc_reference_report:
            assert old['roi_rectangle_xywh']==jsrect,'Fresh reference/browser geometry differs'
            pyrect=old['roi_rectangle_xywh']
        else:
            assert pyrect==jsrect,'Reference/browser rounding changes ROI geometry; rerun corrected reference before using these rows'
        expected_names={'A(弱)','B(强)'} if old['file'].startswith('leica_step') and old['file'] not in ['leica_step14_full.jpg','leica_step15_full.jpg'] else {'整图'}
        assert {r['roiName'] for r in data['perRoi']}==expected_names,'Application output ROI names disagree with harness'
        matches=[r for r in data['perRoi'] if app_geom[r['roiName']]==geom]
        assert len(matches)==1,'Missing/ambiguous geometry match; never fall back to overall'
        matched=matches[0]
        assert old['n']==matched['pixelCount']==pyrect[2]*pyrect[3],'ROI pixel counts disagree'
        pct=100*matched['positiveFraction']
        corrected.append({**old,'original_appPct':historical[(old['file'],old['roi'])]['appPct'],'app_roi_name':matched['roiName'],
            'roi_rectangle_xywh':pyrect,'geometry_source':'historical run_m3.mjs + compare_reference.py CONFIG; no embedded per-run coordinates',
            'appPct':pct,'absDiffPct':abs(pct-old['referencePct']),
            'absDiffAtSameThr':abs(pct-old['referencePctAtAppThr']),
            'mapping_changed':old['roi']!=matched['roiName']})
    runtime_sha='unknown (README declares dd41bae but output JSON has no run-time SHA)'
    if a.app_run_metadata:
        metadata=load(a.app_run_metadata)
        assert metadata['status']=='passed' and metadata['images_failed']==0
        runtime_sha=metadata['source_commit']
        inputs[str(a.app_run_metadata)]=sha(a.app_run_metadata)
    ihc={'audit_type':'Fresh independent reference + fresh pinned browser engine' if a.app_run_metadata else ('Fresh independent reference + historical browser-output geometry rematching; application image engine NOT rerun' if a.ihc_reference_report else 'historical output rematching only; image engines NOT rerun'),
         'image_count':len({r['file'] for r in corrected}),'roi_count':len(corrected),
         'maxAbsDiffPct':max(r['absDiffPct'] for r in corrected),
         'maxAbsDiffAtSameThreshold':max(r['absDiffAtSameThr'] for r in corrected),
         'application_engine_sha':runtime_sha,
         'rows':corrected}
    (a.output/'ihc-corrected-records.json').write_text(json.dumps(ihc,ensure_ascii=False,indent=2))
    pcr=load(pcr_path);wb=load(wb_path)
    claims=[{'claim':'PCR independent-reference checks','count':sum(s['passed']+s['failed'] for s in pcr['scenarios']),
             'passed':sum(s['passed'] for s in pcr['scenarios']),'failed':sum(s['failed'] for s in pcr['scenarios']),
             'evidence_kind':'historical report','runtime_sha':'unknown; README describes original a916051 and subsequent fixed 4142b60'},
            {'claim':'WB 8-bit local-background comparisons','images':sum(i['kind'].startswith('8-bit') for i in wb['images']),
             'lanes':sum(len(i['lanes']) for i in wb['images'] if i['kind'].startswith('8-bit')),
             'max_net_relative_difference':wb['maxNetRelDiff8bit'],'evidence_kind':'historical report',
             'runtime_sha':'unknown; README declares 6bc90ce but no runtime metadata'},
            {'claim':'IHC DAB area comparisons','images':ihc['image_count'],'ROIs':ihc['roi_count'],
             'max_area_difference_percentage_points':ihc['maxAbsDiffPct'],
             'max_difference_at_application_threshold_percentage_points':ihc['maxAbsDiffAtSameThreshold'],
             'evidence_kind':ihc['audit_type'],'runtime_sha':runtime_sha},
            {'claim':'Historical WB synthetic 16-bit','files':2,'measurement_count':12,'unique_constructed_images':1,
             'unique_constructed_lanes':6,'reason':'Generator resets the same seed and uses the same dimensions/parameters for each file.',
             'evidence_kind':'generator inspection; confirm in fresh run output hashes'},
            {'claim':'IHC functional run-through','images':35,'DAB':12,'IF':17,'HE':4,'trichrome':2,
             'limitation':'Functional completion is not independent validation of cell counts, H-score, IF colocalization or biological accuracy.'}]
    (a.output/'claim-evidence.json').write_text(json.dumps(claims,ensure_ascii=False,indent=2))
    (a.output/'input-sha256.json').write_text(json.dumps(inputs,ensure_ascii=False,indent=2))
    print(json.dumps({'IHC_corrected_max_pp':ihc['maxAbsDiffPct'],'IHC_same_threshold_max_pp':ihc['maxAbsDiffAtSameThreshold'],
                      'changed_pairings':sum(r['mapping_changed'] for r in corrected)},indent=2))

if __name__=='__main__':
    main()
```
