## Supplementary Material S1: verification records, scripts and figure source data for "WispLab: open-source browser workbenches for laboratory image and qPCR analysis with reference-based verification": ihc_harness.mjs

```
/**
 * M3 全场景无头管线：dab / he / masson / if 四类真实官网图 → pathology-workbench 官方引擎。
 * 与 M1 相同原理（vite dev + 无头 Chromium 动态 import），按场景选择 analyzeStainImage / analyzeColocImage。
 */
import { createRequire } from 'node:module';
import { readFileSync, writeFileSync, mkdirSync } from 'node:fs';
import { dirname, join } from 'node:path';
import { fileURLToPath } from 'node:url';

const require = createRequire(new URL('./package.json', import.meta.url));
const { chromium } = require('playwright');

const HERE = process.argv[2];
const APP = process.argv[4];
const OUT = process.argv[3];
if (!HERE || !OUT || !APP) throw new Error('Usage: node ihc_harness.mjs IHC_ROOT OUTPUT_DIR VITE_URL');
mkdirSync(OUT, { recursive: true });

const FULL = [{ fx: 0, fy: 0, fw: 1, fh: 1, name: '整图' }];

/** 场景定义：profileId + 文件清单（相对 images/）+ ROI 布局 */
const SCENES = [
  {
    id: 'dab', profile: 'h_dab',
    dir: '', ext: 'jpg',
    files: [
      'leica_step5_full.jpg', 'leica_step6_full.jpg', 'leica_step7_full.jpg',
      'leica_step11_full.jpg', 'leica_step13_full.jpg', 'leica_step14_full.jpg', 'leica_step15_full.jpg',
      'ptglab_ki67_110311.jpg', 'ptglab_ki67_110312.jpg', 'ptglab_ki67_111549.jpg',
      'ptglab_ki67_117470.jpg', 'ptglab_ki67_124444.jpg',
    ],
    roisFor: (f) => (f.startsWith('leica_step') && !['leica_step14_full.jpg', 'leica_step15_full.jpg'].includes(f)
      ? [
          { fx: 0.012, fy: 0.03, fw: 0.46, fh: 0.9, name: 'A(弱)' },
          { fx: 0.53, fy: 0.03, fw: 0.46, fh: 0.9, name: 'B(强)' },
        ]
      : FULL),
  },
  {
    id: 'he', profile: 'h_eosin',
    dir: 'he/', ext: 'jpg',
    files: ['leica_he_fig2.jpg', 'leica_he_fig3.jpg', 'leica_he_fig4.jpg', 'leica_he_fig5.jpg'],
    roisFor: () => FULL,
  },
  {
    id: 'masson', profile: 'masson',
    dir: 'masson/', ext: 'jpg',
    files: ['leica_masson_fig8.jpg', 'leica_gomori_fig16.jpg'],
    roisFor: () => FULL,
  },
  {
    id: 'if', profile: 'if_coloc',
    dir: 'if/', ext: 'jpg',
    files: [
      'leica_if_1000747_001.jpg', 'leica_if_1011587.jpg', 'leica_if_1011589A.jpg', 'leica_if_1012141.jpg',
      'leica_if_1013170B-2.jpg', 'leica_if_1013170B.jpg', 'leica_if_1013184.jpg', 'leica_if_1013205.jpg',
      'leica_if_1013232.jpg', 'leica_if_1013952-A.jpg', 'leica_if_1013952-B.jpg', 'leica_if_1014148A.jpg',
      'leica_if_3-Plex.jpg', 'leica_if_4-Plex.jpg', 'leica_if_5plex_1.jpg',
      'leica_if_CD20-CD3-Tonsil.jpg', 'leica_if_Multiples-staining-2-plex.jpg',
    ],
    roisFor: () => FULL,
  },
];

const browser = await chromium.launch({ args: ['--no-proxy-server'] });
const page = await browser.newPage();
await page.goto(APP, { waitUntil: 'domcontentloaded' });
await page.waitForTimeout(1500);

const summary = [];
for (const scene of SCENES) {
  for (const file of scene.files) {
    const path = join(HERE, 'images', scene.dir, file);
    const b64 = readFileSync(path).toString('base64');
    try {
      const out = await page.evaluate(async ({ b64, name, scene, rois }) => {
        const bin = atob(b64);
        const bytes = new Uint8Array(bin.length);
        for (let i = 0; i < bin.length; i += 1) bytes[i] = bin.charCodeAt(i);
        const { decodeImageFile } = await import('/src/core/image/decode.ts');
        const { analyzeStainImage, analyzeColocImage } = await import('/src/core/analysis/engine.ts');
        const { DEFAULT_THRESHOLD, DEFAULT_SEGMENTATION, DEFAULT_COLOC } = await import('/src/core/types.ts');
        const dec = await decodeImageFile(new File([bytes], name, { type: 'image/jpeg' }));
        const pixels = dec.pixels;
        const W = pixels.width;
        const H = pixels.height;
        let payload;
        if (scene === 'if') {
          const res = analyzeColocImage(pixels, JSON.parse(JSON.stringify(DEFAULT_COLOC)));
          payload = {
            overall: res.overall,
            coloc: res.coloc,
            warnings: res.warnings,
            overlayRgba: res.overlayRgba,
          };
        } else {
          const scaled = rois.map((r, i) => ({
            id: `roi${i}`, name: r.name,
            x: Math.round(r.fx * W), y: Math.round(r.fy * H),
            w: Math.round(r.fw * W), h: Math.round(r.fh * H),
          }));
          const res = analyzeStainImage({
            pixels,
            rois: scaled,
            profileId: scene === 'he' ? 'h_eosin' : scene === 'masson' ? 'masson' : 'h_dab',
            threshold: JSON.parse(JSON.stringify(DEFAULT_THRESHOLD)),
            segmentation: JSON.parse(JSON.stringify(DEFAULT_SEGMENTATION)),
            calibration: { mpp: null },
            stainOverride: null,
          });
          payload = {
            overall: res.overall,
            perRoi: res.perRoi,
            thresholdUsed: res.thresholdUsed,
            nuclearThresholdUsed: res.nuclearThresholdUsed,
            backgroundLevel: res.backgroundLevel,
            warnings: res.warnings,
            qc: res.qc,
            overlayRgba: res.overlayRgba,
          };
        }
        const canvas = document.createElement('canvas');
        canvas.width = W;
        canvas.height = H;
        canvas.getContext('2d').putImageData(new ImageData(new Uint8ClampedArray(payload.overlayRgba), W, H), 0, 0);
        delete payload.overlayRgba;
        return { width: W, height: H, overlay: canvas.toDataURL('image/png'), ...payload };
      }, { b64, name: file, scene: scene.id, rois: scene.roisFor(file) }).catch((e) => { throw new Error(`evaluate: ${e.message}`); });
      const overlayBuf = Buffer.from(out.overlay.split(',')[1], 'base64');
      const outDir = join(OUT, scene.id);
      mkdirSync(outDir, { recursive: true });
      // Vendor-derived overlays are not redistributed; retain numeric outputs only.
      delete out.overlay;
      writeFileSync(join(outDir, file.replace(/\.jpg$/, '.json')), JSON.stringify({ scene: scene.id, file, ...out }, null, 2));
      summary.push({ scene: scene.id, file, ok: true, width: out.width, height: out.height });
      console.log(`✓ ${scene.id}/${file} ${out.width}×${out.height}`);
    } catch (err) {
      summary.push({ scene: scene.id, file, ok: false, error: String(err.message) });
      console.error(`✗ ${scene.id}/${file}: ${err.message}`);
    }
  }
}

writeFileSync(join(OUT, 'summary.json'), JSON.stringify(summary, null, 2));
await browser.close();
const ok = summary.filter((s) => s.ok).length;
console.log(`\n完成：${ok}/${summary.length} 张成功 → ${OUT}`);
```
