## Supplementary Material S1: verification records, scripts and figure source data for "WispLab: open-source browser workbenches for laboratory image and qPCR analysis with reference-based verification": ihc_reference_aligned.py

```
#!/usr/bin/env python3
"""
M3 独立参考实现（与 M1 同策略，不依赖应用代码）：
sips 转 24 位 BMP 解出 RGB → Ruifrok H-DAB 去卷积（独立实现）→ 5% 背景百分位扣除
→ 256 桶 Otsu → 各 ROI 阳性像素占比。与 pathology-workbench 输出逐值比对。
OD 定义 OD255 = 255·ln(255/(I+1))/ln255；独立求逆去卷积；按应用规格对齐端点截断、负浓度截断、float32存储与直方图百分位。
"""
import argparse
import hashlib
import platform
import json
import math
import struct
import subprocess
import sys
import tempfile
from pathlib import Path

parser = argparse.ArgumentParser(description='Rerun independent IHC reference against retained browser outputs. Browser engine is NOT rerun.')
parser.add_argument('--ihc-root', type=Path, required=True)
parser.add_argument('--output', type=Path, required=True)
parser.add_argument('--app-results', type=Path, help='Fresh pinned results_m3 directory; defaults to historical IHC_ROOT/results_m3')
args = parser.parse_args()
HERE = args.ihc_root.resolve()
APP_RESULTS = args.app_results.resolve() if args.app_results else HERE / 'results_m3'
OUT = args.output.resolve()
if OUT.exists():
    raise SystemExit('Output directory must not exist.')
OUT.mkdir(parents=True)
INPUT_HASHES = {}
def hash_input(path):
    INPUT_HASHES[str(path)] = hashlib.sha256(path.read_bytes()).hexdigest()

H_VEC = (0.684, 0.696, 0.183)
DAB_VEC = (0.25, 0.5, 0.85)

def read_bmp_rgb(path):
    """读 24 位 BMP → [R,G,B] 平铺列表（自上而下）。"""
    data = Path(path).read_bytes()
    off = struct.unpack_from('<I', data, 10)[0]
    w = struct.unpack_from('<i', data, 18)[0]
    h = struct.unpack_from('<i', data, 22)[0]
    bpp = struct.unpack_from('<H', data, 28)[0]
    assert bpp == 24, f'BPP {bpp}'
    row_size = (w * 3 + 3) & ~3
    top_down = h < 0  # BMP 高度为负表示自上而下存储
    h_abs = abs(h)
    px = [0.0] * (w * h_abs * 3)
    for row in range(h_abs):
        src = off + row * row_size
        dst_row = row if top_down else h_abs - 1 - row
        base = dst_row * w * 3
        for col in range(w):
            i = src + col * 3
            b, g, r = data[i], data[i + 1], data[i + 2]
            j = base + col * 3
            px[j], px[j + 1], px[j + 2] = r, g, b
    return w, h_abs, px

def stain_od(px, vec, other_vec=H_VEC):
    """标准 Ruifrok 法：OD 向量经染色矩阵求逆投影分离 DAB 通道（255 标尺）。
    与归一化内积捷径不同：求逆法把苏木精串扰从 DAB 通道分离（应用亦用求逆法）。"""
    def norm(v):
        n = math.sqrt(sum(c * c for c in v))
        return [c / n for c in v]
    h = norm(other_vec)
    d = norm(vec)
    res = norm([h[1] * d[2] - h[2] * d[1],
                h[2] * d[0] - h[0] * d[2],
                h[0] * d[1] - h[1] * d[0]])
    # 染色向量作为矩阵的列（与应用 buildStainMatrix 同构），浓度 = inv 第二行 · OD
    m = [[h[0], d[0], res[0]],
         [h[1], d[1], res[1]],
         [h[2], d[2], res[2]]]

    def inv3(mm):
        (a, b, c), (dd, e, f), (g, hh, i) = mm
        det = a * (e * i - f * hh) - b * (dd * i - f * g) + c * (dd * hh - e * g)
        return [[(e * i - f * hh) / det, (c * hh - b * i) / det, (b * f - c * e) / det],
                [(f * g - dd * i) / det, (a * i - c * g) / det, (c * dd - a * f) / det],
                [(dd * hh - e * g) / det, (b * g - a * hh) / det, (a * e - b * dd) / det]]

    inv = inv3(m)
    row = inv[1]
    ln255 = math.log(255.0)
    n_px = len(px) // 3
    out = [0.0] * n_px
    for i in range(n_px):
        r, g, b = px[i * 3] + 1.0, px[i * 3 + 1] + 1.0, px[i * 3 + 2] + 1.0
        ods = [max(0.0, min(255.0, math.log(255.0/v) * 255.0 / ln255)) for v in [r,g,b]]
        od = sum(c*v for c,v in zip(row,ods))
        out[i] = struct.unpack('f',struct.pack('f',max(0.0,od)))[0]
    return out

def otsu_threshold(values):
    hist = [0] * 256
    for v in values:
        b = int(max(0,min(255, math.floor(v+0.5))))
        hist[b] += 1
    total = len(values)
    sum_all = sum(i * hist[i] for i in range(256))
    w_b = sum_b = 0.0
    best = -1.0
    thr = 0
    for i in range(256):
        w_b += hist[i]
        if w_b == 0:
            continue
        w_f = total - w_b
        if w_f == 0:
            break
        sum_b += i * hist[i]
        m_b = sum_b / w_b
        m_f = (sum_all - sum_b) / w_f
        between = w_b * w_f * (m_b - m_f) ** 2
        if between > best:
            best = between
            thr = i
    return thr

CONFIG = {
    'ptglab_ki67_110311.jpg': [{'name': '整图', 'fx': 0, 'fy': 0, 'fw': 1, 'fh': 1}],
    'ptglab_ki67_110312.jpg': [{'name': '整图', 'fx': 0, 'fy': 0, 'fw': 1, 'fh': 1}],
    'ptglab_ki67_111549.jpg': [{'name': '整图', 'fx': 0, 'fy': 0, 'fw': 1, 'fh': 1}],
    'ptglab_ki67_117470.jpg': [{'name': '整图', 'fx': 0, 'fy': 0, 'fw': 1, 'fh': 1}],
    'ptglab_ki67_124444.jpg': [{'name': '整图', 'fx': 0, 'fy': 0, 'fw': 1, 'fh': 1}],
    'leica_step14_full.jpg': [{'name': '整图', 'fx': 0, 'fy': 0, 'fw': 1, 'fh': 1}],
    'leica_step5_full.jpg': [
        {'name': 'A(弱)', 'fx': 0.012, 'fy': 0.03, 'fw': 0.46, 'fh': 0.9},
        {'name': 'B(强)', 'fx': 0.53, 'fy': 0.03, 'fw': 0.46, 'fh': 0.9},
    ],
    'leica_step13_full.jpg': [
        {'name': 'A(深复染)', 'fx': 0.012, 'fy': 0.03, 'fw': 0.46, 'fh': 0.9},
        {'name': 'B(浅复染)', 'fx': 0.53, 'fy': 0.03, 'fw': 0.46, 'fh': 0.9},
    ],
}

def main():
    rows = []
    for fname, rois in CONFIG.items():
        src = HERE / 'images' / fname
        hash_input(src)
        with tempfile.NamedTemporaryFile(suffix='.bmp', delete=False) as tmp:
            bmp = tmp.name
        subprocess.run(['sips', '-s', 'format', 'bmp', str(src), '--out', bmp],
                       check=True, capture_output=True)
        w, h, px = read_bmp_rgb(bmp)
        Path(bmp).unlink()
        dab = stain_od(px, DAB_VEC)
        # 与应用同口径：背景百分位与 Otsu 在整图上计算一次，再套用到各 ROI
        histogram = [0] * 256
        for v in dab:
            histogram[int(max(0,min(255,math.floor(v+0.5))))] += 1
        target = 0.05 * len(dab)
        cumulative = 0
        bg = 255
        for index,count in enumerate(histogram):
            cumulative += count
            if cumulative >= target:
                bg = index
                break
        adj_all = [struct.unpack('f',struct.pack('f',max(0.0,v-bg)))[0] for v in dab]
        thr = otsu_threshold(adj_all)
        app_path = APP_RESULTS / 'dab' / fname.replace('.jpg', '.json')
        hash_input(app_path)
        app_json = json.loads(app_path.read_text())
        app_thr = app_json['thresholdUsed']
        # 参考像素在"应用阈值"下的读数：分离解码器差异与阈值位移两种来源
        ref_at_app = sum(1 for v in adj_all if v > app_thr) / len(adj_all) * 100.0
        row_base = {'referenceThreshold': thr, 'appThreshold': app_thr,
                    'refPositiveAtAppThrWhole': round(ref_at_app, 3)}
        for roi in rois:
            x0 = math.floor(roi['fx'] * w + 0.5)
            y0 = math.floor(roi['fy'] * h + 0.5)
            rw = math.floor(roi['fw'] * w + 0.5)
            rh = math.floor(roi['fh'] * h + 0.5)
            count = 0
            count_app_thr = 0
            n = 0
            for y in range(y0, y0 + rh):
                base = y * w
                for x in range(x0, x0 + rw):
                    n += 1
                    if adj_all[base + x] > thr:
                        count += 1
                    if adj_all[base + x] > app_thr:
                        count_app_thr += 1
            rows.append({'file': fname, 'roi': roi['name'], 'n': n,
                         'referencePct': 100.0 * count / n,
                         'referencePctAtAppThr': 100.0 * count_app_thr / n,
                         'roi_rectangle_xywh': [x0,y0,rw,rh], 'background': round(bg, 2), **row_base})

    # 与应用输出比对
    app_dir = APP_RESULTS / 'dab'
    report = []
    max_abs = 0.0
    max_same_thr = 0.0
    for r in rows:
        data = json.loads((app_dir / (r['file'].replace('.jpg', '.json'))).read_text())
        per = {m['roiName']: m for m in data.get('perRoi', [])}
        # Names differ for step13; both harnesses use the same fractional ROI.
        # Resolve by historical application name after geometry inspection; fail closed.
        expected_name = {'A(深复染)':'A(弱)', 'B(浅复染)':'B(强)'}.get(r['roi'], r['roi'])
        assert expected_name in per, 'Missing application ROI; no overall fallback permitted'
        m = per[expected_name]
        assert m['pixelCount'] == r['n'], 'ROI area mismatch'
        r['application_roi_name'] = expected_name

        app_pct = m['positiveFraction'] * 100
        diff = abs(app_pct - r['referencePct'])
        same_thr_diff = abs(app_pct - r['referencePctAtAppThr'])
        max_abs = max(max_abs, diff)
        max_same_thr = max(max_same_thr, same_thr_diff)
        report.append({**r, 'appPct': round(app_pct, 4), 'absDiffPct': round(diff, 4),
                       'absDiffAtSameThr': round(same_thr_diff, 4)})
    out = {'maxAbsDiffPct': round(max_abs, 4), 'maxAbsDiffAtSameThreshold': round(max_same_thr, 4),
           'rows': report}
    (OUT / 'reference_area_comparison.json').write_text(json.dumps(out, ensure_ascii=False, indent=2))
    metadata = {'operation':'Fresh independent reference from local original images; application outputs are read from APP_RESULTS, with provenance in the separate engine run report.' if args.app_results else 'Fresh independent reference from local original images, compared with retained historical browser outputs; browser engine NOT rerun.',
                'python':sys.version, 'platform':platform.platform(),
                'source_image_and_app_output_sha256':INPUT_HASHES,
                'reference_script_sha256':hashlib.sha256(Path(__file__).read_bytes()).hexdigest(),
                'method_alignment':'Also aligns channel OD endpoint clipping, negative concentration clipping before background, float32 channel storage, nearest-integer histogram bins and percentile background with application specification; no production-code imports.',
                'corrections':['Strict step13 ROI association, no overall fallback','ROI pixel rounding uses floor(x+0.5), matching JavaScript Math.round for these nonnegative coordinates'],
                'application_runtime_sha':'See APP_RESULTS parent run-report.json for separately pinned engine provenance' if args.app_results else 'unknown; historical README declares dd41bae but records do not embed run-time SHA',
                'limitations':['Reference and application image decoders differ','Residual difference is not experimentally attributed to one source; decoders and floating-point evaluation may differ','Same-threshold and separately-thresholded results are both retained']}
    (OUT/'reference-run-metadata.json').write_text(json.dumps(metadata,ensure_ascii=False,indent=2))
    for r in report:
        print(f"{r['file']:<28} {r['roi']:<8} 应用 {r['appPct']:8.3f}%  参考 {r['referencePct']:8.3f}%  |Δ|={r['absDiffPct']:.3f}pp（同阈值 |Δ|={r['absDiffAtSameThr']:.3f}pp，阈值 {r['appThreshold']}/{r['referenceThreshold']}）")
    print(f"最大绝对偏差 {max_abs:.3f}pp；同阈值下最大 {max_same_thr:.3f}pp → qupath/reference_area_comparison.json")

if __name__ == '__main__':
    main()
```
