## Supplementary Material S1: verification records, scripts and figure source data for "WispLab: open-source browser workbenches for laboratory image and qPCR analysis with reference-based verification": pcr_harness.ts

```
/**
 * M1 应用侧提取：调用 pcr-workbench 官方引擎（sniff → extractRows → analyzeDeltaDeltaCt），
 * 把给定场景的完整输入与输出落盘，供独立参考实现逐值比对。
 * 运行：node_modules/.bin/tsx dump_app.ts（在 pcr-workbench 仓库内执行）
 */
import { readFileSync, writeFileSync } from 'node:fs';
import { sniff } from './src/core/sniff';
import { extractRows, analyzeDeltaDeltaCt } from './src/core/analysis';

const OUT_DIR = process.argv[2];
if (!OUT_DIR) throw new Error('Usage: tsx pcr_harness.ts OUTPUT_DIRECTORY');
const matrix: string[][] = JSON.parse(readFileSync(`${OUT_DIR}/matrix.json`, 'utf-8'));

const r = sniff(matrix);
const rows = extractRows(matrix.slice(r.headerRow + 1), r.mapping);

interface Scenario {
  name: string;
  keepSamples?: string[]; // 未指定 = 全部
  refGenes: string[];
  controlSamples: string[];
  efficiencies?: Record<string, number>;
}

const SCENARIOS: Scenario[] = [
  { name: 'S1_livak_welch', keepSamples: ['AiW002-2-D0', 'AiW002-2-D7'], refGenes: ['GAPDH'], controlSamples: ['AiW002-2-D0'] },
  { name: 'S2_livak_anova', refGenes: ['GAPDH', 'ACTB'], controlSamples: ['AiW002-2-D0', 'KYOU-D0'] },
  { name: 'S3_pfaffl', refGenes: ['GAPDH', 'ACTB'], controlSamples: ['AiW002-2-D0', 'KYOU-D0'], efficiencies: { CAMK2A: 92, GRIN1: 105, PAX6: 97, GAPDH: 99, ACTB: 101 } },
];

// S4：中等效应（|t|≈2–3）合成场景，用于刻画 Dunnett 数值缺陷的适用范围。
// Cq 值为手工构造（非真实数据，仅数值路径验证），对照 C1–C3 与处理 T1–T3 各 3 复孔。
const S4_ROWS = (() => {
  const out: { sample: string; target: string; cq: number }[] = [];
  const ref = [19.8, 20.0, 20.2];
  const ctrlGoi: number[][] = [[19.5, 20.0, 20.5], [19.8, 20.2, 20.6], [19.6, 20.1, 20.4]];
  const trtGoi: number[][] = [[20.8, 21.2, 21.6], [20.9, 21.3, 21.7], [20.7, 21.1, 21.5]];
  for (let i = 0; i < 3; i++) {
    for (const v of ref) out.push({ sample: `C${i + 1}`, target: 'REF', cq: v });
    for (const v of ref) out.push({ sample: `T${i + 1}`, target: 'REF', cq: v });
    for (const v of ctrlGoi[i]) out.push({ sample: `C${i + 1}`, target: 'GOI', cq: v });
    for (const v of trtGoi[i]) out.push({ sample: `T${i + 1}`, target: 'GOI', cq: v });
  }
  return out;
})();

for (const sc of SCENARIOS) {
  const kept = sc.keepSamples ? rows.filter((x) => sc.keepSamples!.includes(x.sample)) : rows;
  const out = analyzeDeltaDeltaCt(kept, {
    refGenes: sc.refGenes,
    controlSamples: sc.controlSamples,
    efficiencies: sc.efficiencies,
    qc: { minCq: NaN, maxCq: NaN, absDev: NaN, repSD: NaN },
  });
  writeFileSync(
    `${OUT_DIR}/app_output_${sc.name}.json`,
    JSON.stringify(
      {
        scenario: sc,
        sniff: { headerRow: r.headerRow, mapping: r.mapping, format: r.format },
        inputRows: kept
          .filter((x) => x.sample && x.target && Number.isFinite(x.cq))
          .map((x) => ({ sample: x.sample, target: x.target, cq: x.cq })),
        config: { refGenes: sc.refGenes, controlSamples: sc.controlSamples, efficiencies: sc.efficiencies ?? null },
        effMode: out.effMode,
        refGenesUsed: out.refGenesUsed,
        controlSamples: out.controlSamples,
        warnings: out.warnings,
        ntcReport: out.ntcReport,
        results: out.results,
      },
      null,
      2,
    ),
  );
  console.log(`${sc.name}: ${out.results.length} 条 样本×基因 结果，警告 ${out.warnings.length} 条`);
}

// S4 走合成行
{
  const sc: Scenario = { name: 'S4_moderate_t', refGenes: ['REF'], controlSamples: ['C1', 'C2', 'C3'] };
  const kept = S4_ROWS.map((x, i) => ({
    id: 10_000 + i, plate: '', well: '', sample: x.sample, group: '', pairId: '',
    target: x.target, task: '', cq: x.cq, quantity: NaN, excluded: false,
  }));
  const out = analyzeDeltaDeltaCt(kept, {
    refGenes: sc.refGenes,
    controlSamples: sc.controlSamples,
    qc: { minCq: NaN, maxCq: NaN, absDev: NaN, repSD: NaN },
  });
  writeFileSync(
    `${OUT_DIR}/app_output_${sc.name}.json`,
    JSON.stringify({
      scenario: { ...sc, synthetic: '手工构造 Cq（仅数值路径验证）' },
      sniff: { headerRow: -1, mapping: 'synthetic', format: 'synthetic' },
      inputRows: S4_ROWS,
      config: { refGenes: sc.refGenes, controlSamples: sc.controlSamples, efficiencies: null },
      effMode: out.effMode,
      refGenesUsed: out.refGenesUsed,
      controlSamples: out.controlSamples,
      warnings: out.warnings,
      ntcReport: out.ntcReport,
      results: out.results,
    }, null, 2),
  );
  console.log(`${sc.name}: ${out.results.length} 条 样本×基因 结果，警告 ${out.warnings.length} 条`);
}
```
