## Supplementary Material S1: verification records, scripts and figure source data for "WispLab: open-source browser workbenches for laboratory image and qPCR analysis with reference-based verification": pcr_reference.py

```
#!/usr/bin/env python3
"""
M1 独立参考实现：从应用引擎落盘的输入（孔级 Cq）出发，用独立代码重算
ΔCt / ΔΔCt(Livak) / Pfaffl 效率校正 FC / Welch t / 单因素 ANOVA / Dunnett 校正 p，
与 pcr-workbench 引擎输出逐值比对。不 import 任何仓库代码。

统计数值方法（独立于仓库实现）：
- student t 与 F 分布 p 值经由正则化不完全 beta 函数（math.lgamma + NR 连分式）
- Dunnett 校正 p 经确定性求积重算（同时做 [raw p, k·raw p] 区间校验）
"""
import json
import math
import random
import sys
from pathlib import Path

HERE = Path(sys.argv[1]).resolve() if len(sys.argv) > 1 else Path(__file__).parent
OUT = HERE / 'comparison_report.json'

# ── 数值基础 ──

def mean(xs):
    return sum(xs) / len(xs)

def sd(xs):
    n = len(xs)
    if n < 2:
        return float('nan')
    m = mean(xs)
    return math.sqrt(sum((x - m) ** 2 for x in xs) / (n - 1))

def betacf(a, b, x, max_iter=300, eps=3e-16):
    qab, qap, qam = a + b, a + 1.0, a - 1.0
    c = 1.0
    d = 1.0 - qab * x / qap
    if abs(d) < 1e-300:
        d = 1e-300
    d = 1.0 / d
    h = d
    for m in range(1, max_iter + 1):
        m2 = 2 * m
        aa = m * (b - m) * x / ((qam + m2) * (a + m2))
        d = 1.0 + aa * d
        if abs(d) < 1e-300:
            d = 1e-300
        c = 1.0 + aa / c
        if abs(c) < 1e-300:
            c = 1e-300
        d = 1.0 / d
        h *= d * c
        aa = -(a + m) * (qab + m) * x / ((a + m2) * (qap + m2))
        d = 1.0 + aa * d
        if abs(d) < 1e-300:
            d = 1e-300
        c = 1.0 + aa / c
        if abs(c) < 1e-300:
            c = 1e-300
        d = 1.0 / d
        delta = d * c
        h *= delta
        if abs(delta - 1.0) < eps:
            break
    return h

def betainc_reg(a, b, x):
    """正则化不完全 beta I_x(a,b)。"""
    if x <= 0:
        return 0.0
    if x >= 1:
        return 1.0
    lbeta = math.lgamma(a + b) - math.lgamma(a) - math.lgamma(b)
    front = math.exp(lbeta + a * math.log(x) + b * math.log1p(-x))
    if x < (a + 1.0) / (a + b + 2.0):
        return front * betacf(a, b, x) / a
    return 1.0 - math.exp(lbeta + b * math.log1p(-x) + a * math.log(x)) * betacf(b, a, 1.0 - x) / b

def student_t_two_sided_p(t, df):
    if not math.isfinite(t) or not math.isfinite(df) or df <= 0:
        return float('nan')
    if t == 0:
        return 1.0
    x = df / (df + t * t)
    return betainc_reg(df / 2.0, 0.5, x)

def f_pvalue(f, df1, df2):
    if not math.isfinite(f) or f <= 0:
        return 1.0
    x = df1 * f / (df1 * f + df2)
    return 1.0 - betainc_reg(df1 / 2.0, df2 / 2.0, x)

def normal_cdf(x):
    return 0.5 * (1.0 + math.erf(x / math.sqrt(2.0)))

# ── Dunnett 确定性参考求积（与引擎不同路径：自求 Legendre 节点 + 不完全 Gamma 卡方分位数）──

def gammainc_lower_reg(a, x, iters=500):
    """正则化下不完全 Gamma P(a,x)：级数（x<a+1）与连分式（其余）。"""
    if x <= 0:
        return 0.0
    if x < a + 1.0:
        ap = a
        s = 1.0 / a
        delta = s
        for _ in range(iters):
            ap += 1.0
            delta *= x / ap
            s += delta
            if abs(delta) < abs(s) * 1e-16:
                break
        return s * math.exp(-x + a * math.log(x) - math.lgamma(a))
    # 连分式 Q(a,x) 的 1-Q
    b, c = x + 1.0 - a, 1e300
    d = 1.0 / b
    h = d
    for i in range(1, iters + 1):
        an = -i * (i - a)
        b += 2.0
        d = an * d + b
        if abs(d) < 1e-300:
            d = 1e-300
        c = b + an / c
        if abs(c) < 1e-300:
            c = 1e-300
        d = 1.0 / d
        delta = d * c
        h *= delta
        if abs(delta - 1.0) < 1e-16:
            break
    q = math.exp(-x + a * math.log(x) - math.lgamma(a)) * h
    return 1.0 - q

def chi2_quantile(q, df):
    """卡方分布分位数（二分法反演 P(df/2, x/2)=q）。"""
    lo, hi = 0.0, 1.0
    while gammainc_lower_reg(df / 2.0, hi / 2.0) < q:
        hi *= 2.0
        if hi > 1e12:
            break
    for _ in range(200):
        mid = 0.5 * (lo + hi)
        if gammainc_lower_reg(df / 2.0, mid / 2.0) < q:
            lo = mid
        else:
            hi = mid
    return 0.5 * (lo + hi)

def gl_nodes(n):
    """n 点 Gauss-Legendre 节点与权重（Newton 迭代求 Legendre 根），区间 [-1,1]。"""
    nodes = []
    for i in range(1, n + 1):
        x = math.cos(math.pi * (i - 0.25) / (n + 0.5))
        for _ in range(200):
            p0, p1 = 1.0, x
            for k in range(2, n + 1):
                p0, p1 = p1, ((2 * k - 1) * x * p1 - (k - 1) * p0) / k
            dp = n * (x * p1 - p0) / (x * x - 1.0)
            dx = p1 / dp
            x -= dx
            if abs(dx) < 1e-15:
                break
        nodes.append((x, 2.0 / ((1.0 - x * x) * dp * dp)))
    return nodes

def dunnett_quad(comps, df, n_u=300, panels=60, n_v=24):
    """确定性重算单步 Dunnett 校正 p（χ² 变量域 + 逐比较自适应截断，直接积分尾部）：
    tail = ∫φ(u) ∫₀^vCut Π 差的补 f_χ²ν(v) dv du；vCut 取使窗口亏缺 ≤ 2Φ(−8) 的上界。
    返回 [逐比较校正 p]。"""
    u_nodes = [(-9.0 + 18.0 * (x + 1.0) / 2.0, w * 9.0) for x, w in gl_nodes(n_u)]
    lg = math.lgamma(df / 2.0)
    def chi_pdf(v):
        if v <= 0:
            return 0.0
        return math.exp((df / 2.0 - 1.0) * math.log(v) - v / 2.0 - (df / 2.0) * math.log(2.0) - lg)
    b_max = max(b for _, b in comps)
    out = []
    for t_abs, _ in comps:
        r_min = min(math.sqrt(1.0 + b * b) for _, b in comps)
        s_cut = (8.0 + b_max * 9.0) / (t_abs * r_min)
        v_cut = max(df * s_cut * s_cut, float(df))
        step = v_cut / panels
        half = step / 2.0
        v_nodes = []
        for p in range(panels):
            lo = p * step
            for x, w in gl_nodes(n_v):
                v = lo + half * (x + 1.0)
                v_nodes.append((v, half * w))
        tail = 0.0
        for u, wu in u_nodes:
            inner = 0.0
            for v, wv in v_nodes:
                s = math.sqrt(v / df)
                prod = 1.0
                for b in (c[1] for c in comps):
                    r = math.sqrt(1.0 + b * b)
                    a = s * t_abs * r
                    off = b * u
                    prod *= normal_cdf(a + off) - normal_cdf(-a + off)
                    if prod == 0.0:
                        break
                inner += wv * chi_pdf(v) * (1.0 - prod)
            tail += wu * (math.exp(-0.5 * u * u) / math.sqrt(2.0 * math.pi)) * inner
        out.append(max(0.0, min(1.0, tail)))
    return out

def self_check():
    """对照统计表验证数值实现，偏差过大直接终止。"""
    p10 = student_t_two_sided_p(2.0, 10.0)
    assert abs(p10 - 0.07338803) < 1e-6, f't(2.0, df=10) 双侧 p {p10}'
    p1 = student_t_two_sided_p(1.0, 1.0)
    assert abs(p1 - 0.5) < 1e-9, f't(1, df=1) 双侧 p 应为 0.5，得到 {p1}'
    pf = f_pvalue(4.9646, 2, 10)  # 临界值对应 p≈0.0314
    assert 0.028 < pf < 0.035, f'F(4.9646, 2,10) p {pf}'

# ── 参考实现 ──

def analyze(rows, ref_genes, control_samples, efficiencies):
    """rows: [{sample,target,cq}]，已排除 NTC/标准品（本 fixture 无）。"""
    eff_pct = lambda g: (efficiencies or {}).get(g, 100)
    # 1. 复孔聚合
    agg = {}
    for r in rows:
        agg.setdefault((r['sample'], r['target']), []).append(r['cq'])
    samples = sorted({r['sample'] for r in rows})
    genes = sorted({r['target'] for r in rows})
    refs = [g for g in ref_genes if g in genes]
    targets = [g for g in genes if g not in refs]
    controls = [s for s in control_samples if s in samples]
    mean_cq = {k: mean(v) for k, v in agg.items()}

    # 2. 内参池化（每样本取各内参 meanCq 的均值）
    ref_cq = {}
    for s in samples:
        vals = [mean_cq[(s, g)] for g in refs if (s, g) in mean_cq]
        if vals:
            ref_cq[s] = mean(vals)

    # 3. 个体 ΔCt
    delta = {}
    for g in targets:
        for s in samples:
            if (s, g) not in agg or s not in ref_cq:
                continue
            vals = agg[(s, g)]
            dvals = [v - ref_cq[s] for v in vals]
            delta[(s, g)] = {'values': vals, 'ref_cq': ref_cq[s], 'dvals': dvals,
                             'delta_ct': mean(dvals)}

    report = {'ref_genes_used': refs, 'controls': controls, 'results': {}}
    for g in targets:
        pooled_ctrl_d = []
        for s in controls:
            pooled_ctrl_d.extend(delta[(s, g)]['dvals'])
        if not pooled_ctrl_d:
            continue
        ctrl_mean = mean(pooled_ctrl_d)
        pooled_ctrl_raw = [v for s in controls for v in delta[(s, g)]['values']]
        ct_t_ctrl = mean(pooled_ctrl_raw)
        ct_r_ctrl = mean([delta[(s, g)]['ref_cq'] for s in controls])
        e_t = 1 + eff_pct(g) / 100
        e_r = mean([1 + eff_pct(x) / 100 for x in refs]) if refs else 2.0
        eff_active = abs(e_t - 2) > 1e-9 or abs(e_r - 2) > 1e-9
        gene_eff_mode = eff_active

        # 统计输入
        entries = [(s, delta[(s, g)]) for s in samples if (s, g) in delta]
        ctrl_vals = pooled_ctrl_d
        use_anova = len(entries) > 2 and len(samples) > 2 and len(ctrl_vals) >= 2
        anova = None
        if use_anova:
            groups = []
            for s, d in entries:
                if s in controls or len(d['dvals']) < 2:
                    continue
                groups.append(list(d['dvals']))
            if len(groups) >= 2:
                allv = [v for grp in groups for v in grp]
                grand = mean(allv)
                ssb = sum(len(grp) * (mean(grp) - grand) ** 2 for grp in groups)
                ssw = sum((x - mean(grp)) ** 2 for grp in groups for x in grp)
                k, n = len(groups), len(allv)
                df1, df2 = k - 1, n - k
                msw = ssw / df2
                f_stat = (ssb / df1) / msw if msw > 0 else 0.0
                anova = {'f': f_stat, 'df1': df1, 'df2': df2, 'p': f_pvalue(f_stat, df1, df2), 'ms_within': msw}

        for s, d in entries:
            is_ctrl = s in controls
            if gene_eff_mode:
                fc_values = [e_t ** (ct_t_ctrl - v) / e_r ** (ct_r_ctrl - d['ref_cq']) for v in d['values']]
                fold_change = mean(fc_values)
            else:
                fc_values = [2 ** -(v - ctrl_mean) for v in d['dvals']]
                fold_change = 2 ** -(d['delta_ct'] - ctrl_mean)
            p_value = p_adj = None
            method = ''
            if not is_ctrl and len(ctrl_vals) >= 2:
                if use_anova and anova and math.isfinite(anova['p']):
                    p_value = anova['p']
                    method = 'ANOVA'
                if p_value is None:
                    method = 'Welch'
                if method == 'Welch':
                    m1, m2 = mean(d['dvals']), mean(ctrl_vals)
                    v1, v2 = sd(d['dvals']) ** 2, sd(ctrl_vals) ** 2
                    se = math.sqrt(v1 / len(d['dvals']) + v2 / len(ctrl_vals))
                    if not math.isfinite(se) or se == 0:
                        t_stat, df_w, p_value = 0.0, len(d['dvals']) + len(ctrl_vals) - 2, 1.0
                    else:
                        t_stat = (m1 - m2) / se
                        num = (v1 / len(d['dvals']) + v2 / len(ctrl_vals)) ** 2
                        den = (v1 / len(d['dvals'])) ** 2 / (len(d['dvals']) - 1) + (v2 / len(ctrl_vals)) ** 2 / (len(ctrl_vals) - 1)
                        df_w = num / den if den > 0 else len(d['dvals']) + len(ctrl_vals) - 2
                        p_value = student_t_two_sided_p(abs(t_stat), df_w)
            report['results'].setdefault(g, {})[s] = {
                'is_control': is_ctrl,
                'n': len(d['values']),
                'mean_cq': mean(d['values']),
                'sd_cq': sd(d['values']),
                'ref_cq': d['ref_cq'],
                'delta_ct': d['delta_ct'],
                'dd_ct': d['delta_ct'] - ctrl_mean,
                'ctrl_delta_mean': ctrl_mean,
                'eff_mode': gene_eff_mode,
                'fold_change': fold_change,
                'fc_mean': mean(fc_values),
                'fc_sd': sd(fc_values),
                'fc_values': fc_values,
                'stat_method': method if not is_ctrl else '',
                'welch': ({'t': t_stat, 'df': df_w, 'p': p_value} if method == 'Welch' else None),
                'anova': anova,
                'p_value': p_value,
            }
            if use_anova and not is_ctrl:
                dvals = delta[(s, g)]['dvals']
                report['results'][g][s]['dunnett_t'] = abs(mean(dvals) - ctrl_mean) / (
                    math.sqrt(anova['ms_within']) * math.sqrt(1 / len(dvals) + 1 / len(ctrl_vals)))
                report['results'][g][s]['dunnett_b'] = math.sqrt(len(dvals) / len(ctrl_vals))
                report['results'][g][s]['anova_df2'] = anova['df2']
    return report

def dunnett_mc_multi(comps, df, n_draws=400_000, seed=20261017):
    """固定种子蒙特卡洛重算 Dunnett 单步校正 p（联合通过概率口径）：
    adj = 1 − P(对所有比较 |Z_i + b_i·u| ≤ c_i·s·√(1+b_i²))，u~N(0,1)，s=√(χ²_df/df)。"""
    rng = random.Random(seed)
    all_ok = 0
    for _ in range(n_draws):
        u = rng.gauss(0, 1)
        chi2 = sum(rng.gauss(0, 1) ** 2 for _ in range(df))
        s = math.sqrt(chi2 / df)
        ok = all(abs(rng.gauss(0, 1) + b * u) <= t_abs * s * math.sqrt(1 + b * b)
                 for t_abs, b in comps)
        all_ok += ok
    return [1.0 - all_ok / n_draws] * len(comps), n_draws

def stars(p):
    if p is None or not math.isfinite(p):
        return ''
    if p < 0.001:
        return '***'
    if p < 0.01:
        return '**'
    if p < 0.05:
        return '*'
    return 'ns'

def rel_diff(a, b):
    scale = max(abs(a), abs(b), 1e-12)
    return abs(a - b) / scale

def compare():
    self_check()
    summary = {'self_check': 'ok', 'scenarios': []}
    all_pass = True
    for path in sorted(HERE.glob('app_output_*.json')):
        data = json.loads(path.read_text())
        name = data['scenario']['name']
        ref = analyze(data['inputRows'], data['config']['refGenes'],
                      data['config']['controlSamples'], data['config']['efficiencies'])
        checks = []
        n_pass = n_fail = 0

        def check(field, app_v, ref_v, tol, kind='abs'):
            nonlocal n_pass, n_fail, all_pass
            if app_v is None and ref_v is None:
                checks.append({'field': field, 'status': 'both-null'})
                return
            if kind == 'rel':
                diff = rel_diff(app_v, ref_v)
            else:
                diff = abs((app_v or 0) - (ref_v or 0))
            ok = diff <= tol
            n_pass, n_fail = n_pass + ok, n_fail + (not ok)
            all_pass = all_pass and ok
            checks.append({'field': field, 'app': app_v, 'ref': ref_v,
                           'diff': diff, 'tol': tol, 'ok': ok})

        dunnett_pool = []
        for g, samples in ref['results'].items():
            for s, rr in samples.items():
                app = next(x for x in data['results'] if x['target'] == g and x['sample'] == s)
                key = f'{g}/{s}'
                check(f'{key}.n', app['n'], rr['n'], 0)
                check(f'{key}.meanCq', app['meanCq'], rr['mean_cq'], 1e-9)
                check(f'{key}.refCq', app['refCq'], rr['ref_cq'], 1e-9)
                check(f'{key}.deltaCt', app['deltaCt'], rr['delta_ct'], 1e-9)
                check(f'{key}.ddCt', app['ddCt'], rr['dd_ct'], 1e-9)
                check(f'{key}.fcMean', app['fcMean'], rr['fc_mean'], 1e-9, 'rel')
                check(f'{key}.fcSd', app['fcSd'] if math.isfinite(app['fcSd']) else None,
                      rr['fc_sd'] if math.isfinite(rr['fc_sd']) else None, 1e-9, 'rel')
                check(f'{key}.foldChange', app['foldChange'], rr['fold_change'], 1e-9, 'rel')
                # 效率模式判定一致
                ok_eff = app_is_eff = ('Pfaffl' in (app['statMethod'] or '')) == rr['eff_mode']
                if rr['is_control']:
                    ok_eff = True
                n_pass, n_fail = n_pass + ok_eff, n_fail + (not ok_eff)
                all_pass = all_pass and ok_eff
                checks.append({'field': f'{key}.effMode', 'app': app['statMethod'], 'ref': rr['eff_mode'], 'ok': ok_eff})
                if rr['stat_method'] == 'Welch':
                    # statMethod 字符串仅保留 2 位小数（t.toFixed(2)），故 t 容差取 0.0051；p 值为完整精度比对
                    check(f'{key}.welch.t(str2dp)', app_t(app), rr['welch']['t'], 0.0051)
                    check(f'{key}.welch.p', app['pValue'], rr['welch']['p'], 1e-8)
                elif rr['stat_method'] == 'ANOVA':
                    check(f'{key}.anova.p', app['pValue'], rr['anova']['p'], 1e-8)
                    if 'dunnett_t' in rr:
                        dunnett_pool.append({'key': key, 'app_p': app['pAdjusted'], 'ref': rr})
                else:
                    ok_np = app['pValue'] is None
                    n_pass, n_fail = n_pass + ok_np, n_fail + (not ok_np)
                    all_pass = all_pass and ok_np
                    checks.append({'field': f'{key}.pValue', 'app': app['pValue'], 'ref': None, 'ok': ok_np})
                # 显著性星标与参考 p 一致
                ref_p_for_stars = None
                if rr['stat_method'] == 'Welch':
                    ref_p_for_stars = rr['welch']['p']
                elif rr['stat_method'] == 'ANOVA':
                    ref_p_for_stars = rr['anova']['p']
                if ref_p_for_stars is not None:
                    expected = stars(app['pAdjusted']) if app['pAdjusted'] is not None else stars(ref_p_for_stars)
                    ok_stars = expected == app['significance'] if not rr['is_control'] else app['significance'] == ''
                    n_pass, n_fail = n_pass + ok_stars, n_fail + (not ok_stars)
                    all_pass = all_pass and ok_stars
                    checks.append({'field': f'{key}.stars', 'app': app['significance'], 'ref': expected, 'ok': ok_stars})

        # Dunnett 校正 p：确定性双维 Gauss-Legendre 重算（300×200 节点，独立于引擎的 40×40）
        # 区间校验用逐比较 raw p（t_i, df2）：单步校正 p 满足 raw ≤ adj ≤ min(1, k·raw)
        by_gene = {}
        for item in dunnett_pool:
            by_gene.setdefault(item['key'].split('/')[0], []).append(item)
        for g, items in by_gene.items():
            comps = [(it['ref']['dunnett_t'], it['ref']['dunnett_b']) for it in items]
            df2 = items[0]['ref']['anova_df2']
            quad_ps = dunnett_quad(comps, df2)
            for it, quad_p in zip(items, quad_ps):
                raw_p = student_t_two_sided_p(it['ref']['dunnett_t'], df2)
                lo, hi = raw_p, min(1.0, raw_p * len(comps))
                ok_iv = lo - 1e-12 <= it['app_p'] <= hi + 1e-12
                rel = abs(it['app_p'] - quad_p) / max(abs(it['app_p']), abs(quad_p), 1e-300)
                ok_quad = rel <= 0.02
                n_pass, n_fail = n_pass + (ok_iv and ok_quad), n_fail + (not (ok_iv and ok_quad))
                all_pass = all_pass and ok_iv and ok_quad
                checks.append({'field': f"{it['key']}.dunnettP", 'app': it['app_p'], 'ref_quad': quad_p,
                               'rel_diff': rel, 'raw_p_interval': [lo, hi],
                               'ok': ok_iv and ok_quad})

        # 警告与结构
        ok_warn = data['warnings'] == []
        checks.append({'field': 'warnings-empty', 'app': data['warnings'], 'ok': ok_warn})
        n_pass, n_fail = n_pass + ok_warn, n_fail + (not ok_warn)
        all_pass = all_pass and ok_warn

        summary['scenarios'].append({
            'name': name, 'passed': n_pass, 'failed': n_fail, 'all_pass': n_fail == 0,
            'checks': checks,
        })
    OUT.write_text(json.dumps(summary, ensure_ascii=False, indent=2))
    for sc in summary['scenarios']:
        flag = 'PASS' if sc['all_pass'] else 'FAIL'
        print(f"[{flag}] {sc['name']}: {sc['passed']} 项一致, {sc['failed']} 项不一致")
    print(f"报告: {OUT}")
    sys.exit(0 if all(s['all_pass'] for s in summary['scenarios']) else 1)

def app_t(app):
    m = app.get('statMethod') or ''
    if 't=' in m:
        return float(m.split('t=')[1].split(',')[0])
    return None

def dunnett_mc_multi(comps, df, n_draws=400_000, seed=20261017):
    """（已弃用，保留说明）蒙特卡洛在 p ≲ 1e-5 量级分辨率不足，已由 dunnett_quad 确定性求积取代。"""
    raise NotImplementedError('由 dunnett_quad 取代')

if __name__ == '__main__':
    compare()
```
