## Supplementary Material S1: verification records, scripts and figure source data for "WispLab: open-source browser workbenches for laboratory image and qPCR analysis with reference-based verification": prepare_public_copy.py

```
#!/usr/bin/env python3
"""Create an allowlisted, text-only, de-identified LOCAL copy; never publish."""
import argparse
import hashlib
import json
from pathlib import Path
import re

ROOT=Path(__file__).resolve().parent.parent

def main():
    parser=argparse.ArgumentParser()
    parser.add_argument('--output',type=Path,required=True)
    args=parser.parse_args()
    out=args.output.resolve()
    if out.exists():
        raise SystemExit('Output directory must not already exist.')
    out.mkdir(parents=True)
    allow=json.loads((ROOT/'archive-allowlist.json').read_text())
    hashes={}
    for name in allow['files']:
        rel=Path(name)
        if rel.is_absolute() or '..' in rel.parts:
            raise ValueError('Unsafe relative path')
        source=ROOT/rel
        if source.suffix.lower() not in {'.md','.py','.ts','.mjs','.json','.csv','.tsv','.log','.txt'}:
            raise ValueError('Non-text payload rejected')
        text=source.read_text().replace(str(Path.home()),'<LOCAL_HOME>')
        text=re.sub(r'(?:/private)?/var/folders/[^/\s]+/[^/\s]+/T/[^/\s]+', '<LOCAL_TEMP>', text)
        target=out/rel
        target.parent.mkdir(parents=True,exist_ok=True)
        target.write_text(text)
        hashes[name]=hashlib.sha256(target.read_bytes()).hexdigest()
    manifest={'operation':'Local allowlisted copy only; not published',
              'transformation':'Home-directory prefix replaced with <LOCAL_HOME>; macOS temporary-directory prefix replaced with <LOCAL_TEMP>; original evidence unchanged',
              'files_sha256':hashes}
    (out/'PUBLIC-COPY-MANIFEST.json').write_text(json.dumps(manifest,indent=2))
    print(f'Prepared {len(hashes)} text files in {out}')

if __name__=='__main__':
    main()
```
