## Supplementary Material S1: verification records, scripts and figure source data for "WispLab: open-source browser workbenches for laboratory image and qPCR analysis with reference-based verification": run_ihc_pinned.py

```
#!/usr/bin/env python3
"""Run historical 35-image harness using isolated pinned pathology source.

Requires locally acquired images and existing Playwright browser installation.
Does not download or save source images/overlays into the output directory.
"""
import argparse
import hashlib
import json
from pathlib import Path
import shutil
import socket
import subprocess
import tempfile
import time
import urllib.request
from datetime import datetime,timezone
from run_reproduction import extract,sha,cmd,SCRIPTS

PIN='dd41bae4bc27fda0b6bf1135b802e4af41ae7316'

def main():
    ap=argparse.ArgumentParser()
    ap.add_argument('--repo',type=Path,required=True)
    ap.add_argument('--ihc-root',type=Path,required=True)
    ap.add_argument('--output',type=Path,required=True)
    ap.add_argument('--node-modules',type=Path)
    a=ap.parse_args()
    out=a.output.resolve()
    if out.exists():
        raise SystemExit('Output directory must not already exist.')
    out.mkdir(parents=True)
    deps=(a.node_modules or a.repo/'node_modules').resolve()
    report={'source_commit':PIN,'started_utc':datetime.now(timezone.utc).isoformat(),
            'operation':'Fresh browser-engine run from isolated pinned source; no UI clicks or biological validation.',
            'dependency_install':'Existing node_modules and Playwright browser reused.',
            'status':'failed','node':cmd(['node','--version']),
            'packages':{p:json.loads((deps/p/'package.json').read_text())['version'] for p in ['vite','playwright','typescript']}}
    proc=None
    try:
        with tempfile.TemporaryDirectory(prefix='wisplab-ihc-source-') as tmp:
            stage=Path(tmp)
            extract(a.repo.resolve(),PIN,stage)
            report['source_file_sha256']={str(p.relative_to(stage)):sha(p) for p in stage.rglob('*') if p.is_file()}
            report['harness_sha256']=sha(SCRIPTS/'ihc_harness.mjs')
            shutil.copy2(SCRIPTS/'ihc_harness.mjs',stage/'ihc_harness.mjs')
            (stage/'node_modules').symlink_to(deps,target_is_directory=True)
            with socket.socket() as sock:
                sock.bind(('127.0.0.1',0))
                port=sock.getsockname()[1]
            url='http://127.0.0.1:'+str(port)
            with (out/'vite.log').open('w') as log:
                proc=subprocess.Popen([str(deps/'.bin/vite'),'--host','127.0.0.1','--port',str(port),'--strictPort'],cwd=stage,stdout=log,stderr=log)
                for attempt in range(60):
                    try:
                        with urllib.request.urlopen(url,timeout=1) as response:
                            assert response.status==200
                        break
                    except Exception:
                        if proc.poll() is not None:
                            raise RuntimeError('Vite exited; see log')
                        time.sleep(0.5)
                else:
                    raise RuntimeError('Vite did not start')
                run=subprocess.run(['node',str(stage/'ihc_harness.mjs'),str(a.ihc_root.resolve()),str(out/'results_m3'),url],cwd=stage,capture_output=True,text=True,timeout=300)
                (out/'browser.stdout.log').write_text(run.stdout)
                (out/'browser.stderr.log').write_text(run.stderr)
                report['browser_exit_code']=run.returncode
                if run.returncode:
                    raise RuntimeError('Browser harness failed')
                summary=json.loads((out/'results_m3/summary.json').read_text())
                report['summary']=summary
                report['images_completed']=sum(r['ok'] for r in summary)
                report['images_failed']=sum(not r['ok'] for r in summary)
                report['input_sha256']={str(p.relative_to(a.ihc_root)):sha(p) for p in (a.ihc_root/'images').rglob('*') if p.is_file()}
                if report['images_failed']:
                    raise RuntimeError('At least one image failed')
                report['status']='passed'
    except Exception as exc:
        report['error']=repr(exc)
    finally:
        if proc is not None:
            proc.terminate()
            try:
                proc.wait(timeout=10)
            except subprocess.TimeoutExpired:
                proc.kill()
        report['finished_utc']=datetime.now(timezone.utc).isoformat()
        (out/'run-report.json').write_text(json.dumps(report,ensure_ascii=False,indent=2))
    print(json.dumps({k:v for k,v in report.items() if k in ['status','images_completed','images_failed','error']},ensure_ascii=False))
    return 0 if report['status']=='passed' else 1

if __name__=='__main__':
    raise SystemExit(main())
```
