## Supplementary Material S1: verification records, scripts and figure source data for "WispLab: open-source browser workbenches for laboratory image and qPCR analysis with reference-based verification": run_reproduction.py

```
#!/usr/bin/env python3
"""Rerun pinned WispLab engines without changing any production checkout.

Existing installed node_modules are reused; this is source-isolated, not a
clean dependency installation. The report records this explicitly.
"""
import argparse
import csv
import hashlib
import io
import json
import os
from pathlib import Path
import platform
import shutil
import subprocess
import sys
import tarfile
import tempfile
from datetime import datetime, timezone

PINS = {'pcr': '4142b6078251d3973667bb59352a016ac0ebf3da',
        'wb': '6bc90cec4422d3e414629ce3d78b639c75279588'}
SCRIPTS = Path(__file__).resolve().parent

def sha(path):
    return hashlib.sha256(Path(path).read_bytes()).hexdigest()

def cmd(args, cwd=None):
    return subprocess.run([str(x) for x in args], cwd=cwd, text=True,
                          capture_output=True, check=True).stdout.strip()

def extract(repo, pin, dst):
    actual = cmd(['git', '-C', repo, 'rev-parse', pin + '^{commit}'])
    if actual != pin:
        raise ValueError('Pin did not resolve exactly')
    data = subprocess.run(['git', '-C', str(repo), 'archive', pin],
                          capture_output=True, check=True).stdout
    with tarfile.open(fileobj=io.BytesIO(data)) as tar:
        for member in tar.getmembers():
            if member.name.startswith('/') or '..' in Path(member.name).parts:
                raise ValueError('Unsafe archive member')
        tar.extractall(dst)
    return actual

def main():
    ap = argparse.ArgumentParser()
    ap.add_argument('--pcr-repo', type=Path, required=True)
    ap.add_argument('--wb-repo', type=Path, required=True)
    ap.add_argument('--pcr-node-modules', type=Path)
    ap.add_argument('--wb-node-modules', type=Path)
    ap.add_argument('--output', type=Path, required=True)
    args = ap.parse_args()
    out = args.output.resolve()
    if out.exists():
        raise SystemExit('Output directory must not already exist (preserves prior runs).')
    out.mkdir(parents=True)
    report = {'started_utc': datetime.now(timezone.utc).isoformat(),
              'environment': {'platform': platform.platform(), 'python': sys.version,
                              'node': cmd(['node', '--version'])},
              'dependency_install': 'Existing node_modules reused, no clean npm install performed.',
              'engine_runs': [], 'not_rerun': ['IHC browser engine', 'WB vendor-image engine',
                    'ImageJ end-to-end', 'QuPath', 'IF colocalization validation', 'cell counts/H-score validation']}
    with tempfile.TemporaryDirectory(prefix='wisplab-paper-repro-') as tmp:
        for module in ['pcr', 'wb']:
            repo = getattr(args, module + '_repo').resolve()
            deps = (getattr(args, module + '_node_modules') or repo / 'node_modules').resolve()
            stage = Path(tmp) / module
            stage.mkdir()
            result = {'module': module, 'git_commit': PINS[module], 'status': 'failed',
                      'source_repository': str(repo), 'dependency_directory': str(deps)}
            report['engine_runs'].append(result)
            dst = out / module
            dst.mkdir()
            try:
                extract(repo, PINS[module], stage)
                result['tracked_source_file_count'] = len([p for p in stage.rglob('*') if p.is_file()])
                result['source_file_sha256'] = {str(p.relative_to(stage)):sha(p) for p in stage.rglob('*') if p.is_file()}
                (stage / 'node_modules').symlink_to(deps, target_is_directory=True)
                result['pinned_package_lock_sha256'] = sha(stage / 'package-lock.json')
                result['installed_lock_sha256'] = sha(deps / '.package-lock.json') if (deps / '.package-lock.json').exists() else None
                packages = ['tsx', 'typescript', 'fast-png', 'tiff', 'fflate']
                result['installed_packages'] = {p:json.loads((deps/p/'package.json').read_text())['version'] for p in packages if (deps/p/'package.json').exists()}
                harness = SCRIPTS / ('pcr_harness.ts' if module == 'pcr' else 'wb_synthetic.ts')
                harness_dst = stage / harness.name
                shutil.copy2(harness, harness_dst)
                result['harness_sha256'] = sha(harness)
                if module == 'pcr':
                    fixture = stage / 'tests/fixtures/real/RELATIVE_example.csv'
                    result['fixture_sha256'] = sha(fixture)
                    rows = list(csv.reader(fixture.open(newline='', encoding='utf-8-sig')))
                    rows = [r for r in rows if any(c.strip() for c in r)]
                    (dst/'matrix.json').write_text(json.dumps(rows,ensure_ascii=False))
                invocation = [deps/'.bin/tsx', harness_dst, dst]
                result['invocation_shape'] = ['NODE_MODULES/.bin/tsx', 'EXTRACTED_SOURCE/'+harness.name, 'OUTPUT/'+module]
                run = subprocess.run([str(x) for x in invocation], cwd=stage, capture_output=True, text=True, timeout=180)
                (dst/'engine.stdout.log').write_text(run.stdout)
                (dst/'engine.stderr.log').write_text(run.stderr)
                result['engine_exit_code'] = run.returncode
                if run.returncode:
                    raise RuntimeError('Engine failed; see stderr log')
                if module == 'pcr':
                    reference = SCRIPTS/'pcr_reference.py'
                    result['reference_sha256'] = sha(reference)
                    ref = subprocess.run([sys.executable, str(reference), str(dst)], capture_output=True, text=True, timeout=600)
                    (dst/'reference.stdout.log').write_text(ref.stdout)
                    (dst/'reference.stderr.log').write_text(ref.stderr)
                    result['reference_exit_code'] = ref.returncode
                    if ref.returncode:
                        raise RuntimeError('Independent reference check failed')
                    checks=json.loads((dst/'comparison_report.json').read_text())
                    result['scenarios']=[{k:s[k] for k in ['name','passed','failed','all_pass']} for s in checks['scenarios']]
                    result['checks_passed']=sum(s['passed'] for s in checks['scenarios'])
                    result['checks_failed']=sum(s['failed'] for s in checks['scenarios'])
                else:
                    truth=json.loads((dst/'truth_16.json').read_text())
                    app=json.loads((dst/'app_results.json').read_text())
                    differences=[]
                    for actual,expected in zip(app,truth):
                        if actual['bitDepth'] != 16:
                            raise RuntimeError('Not a decoded 16-bit raster')
                        for a,e in zip(actual['lanes'],expected['lanes']):
                            differences.append({'file':actual['file'],'lane':a['lane'],
                                                'app_net':a['net'],'constructed_net':e['net'],
                                                'absolute_difference':abs(a['net']-e['net'])})
                    maxdiff=max(d['absolute_difference'] for d in differences)
                    file_hashes={p.name:sha(p) for p in dst.glob('*.tif')}
                    wb_report={'measurements':differences,'maximum_absolute_difference':maxdiff,
                               'synthetic_file_sha256':file_hashes,'unique_synthetic_images':len(set(file_hashes.values())),
                               'measurement_count':len(differences),'unique_constructed_lanes':6,
                               'interpretation':'Two identical generated TIFF repetitions; not two independent test images.'}
                    (dst/'comparison_report.json').write_text(json.dumps(wb_report,indent=2))
                    result.update({k:v for k,v in wb_report.items() if k!='measurements'})
                    if maxdiff != 0:
                        raise RuntimeError('Synthetic measurement differs from constructed truth')
                result['status']='passed'
            except Exception as exc:
                result['error']=repr(exc)
            result['output_sha256']={str(p.relative_to(dst)):sha(p) for p in dst.rglob('*') if p.is_file()}
            (out/'run-report.json').write_text(json.dumps(report,ensure_ascii=False,indent=2))
    report['finished_utc']=datetime.now(timezone.utc).isoformat()
    report['all_requested_runs_passed']=all(r['status']=='passed' for r in report['engine_runs'])
    (out/'run-report.json').write_text(json.dumps(report,ensure_ascii=False,indent=2))
    print(json.dumps([{k:v for k,v in r.items() if k in ['module','status','error','checks_passed','checks_failed','maximum_absolute_difference']} for r in report['engine_runs']],indent=2))
    return 0 if report['all_requested_runs_passed'] else 1

if __name__=='__main__':
    raise SystemExit(main())
```
