## Supplementary Material S1: verification records, scripts and figure source data for "WispLab: open-source browser workbenches for laboratory image and qPCR analysis with reference-based verification": run_wb_real_pinned.py

```
#!/usr/bin/env python3
"""Pinned WB image-engine + independent reference rerun; retains numbers only."""
import argparse
import csv
import json
import os
from pathlib import Path
import shutil
import subprocess
import sys
import tempfile
from datetime import datetime,timezone
from run_reproduction import PINS,extract,sha,SCRIPTS,cmd

def main():
    ap=argparse.ArgumentParser()
    ap.add_argument('--repo',type=Path,required=True)
    ap.add_argument('--corpus',type=Path,required=True)
    ap.add_argument('--output',type=Path,required=True)
    a=ap.parse_args()
    out=a.output.resolve()
    if out.exists():
        raise SystemExit('Output directory must not already exist.')
    out.mkdir(parents=True)
    deps=(a.repo/'node_modules').resolve()
    report={'git_commit':PINS['wb'],'started_utc':datetime.now(timezone.utc).isoformat(),
            'status':'failed','operation':'Fresh pinned decode/proposal/local-background measurement plus independent Python reference; no ImageJ.',
            'dependency_install':'Existing node_modules reused. Image conversion uses macOS sips.',
            'node':cmd(['node','--version']),'source_corpus_manifest_sha256':sha(a.corpus/'MANIFEST.json')}
    try:
        with tempfile.TemporaryDirectory(prefix='wisplab-wb-real-') as tmp:
            stage=Path(tmp)/'source';stage.mkdir()
            data=Path(tmp)/'data';data.mkdir()
            extract(a.repo.resolve(),PINS['wb'],stage)
            report['source_file_sha256']={str(p.relative_to(stage)):sha(p) for p in stage.rglob('*') if p.is_file()}
            (stage/'node_modules').symlink_to(deps,target_is_directory=True)
            harness=SCRIPTS/'wb_real_harness.ts'
            shutil.copy2(harness,stage/harness.name)
            report['harness_sha256']=sha(harness)
            run=subprocess.run([str(deps/'.bin/tsx'),str(stage/harness.name),str(a.corpus.resolve()),str(data)],cwd=stage,capture_output=True,text=True,timeout=180)
            (out/'engine.stdout.log').write_text(run.stdout)
            (out/'engine.stderr.log').write_text(run.stderr)
            if run.returncode:
                raise RuntimeError('Engine failed')
            report['converted_input_sha256']={p.name:sha(p) for p in (data/'staged').glob('*') if p.is_file()}
            report['original_input_sha256']={p.name:sha(p) for p in (data/'inputs').glob('*') if p.is_file()}
            ref=subprocess.run([sys.executable,str(SCRIPTS/'wb_reference.py'),str(data)],capture_output=True,text=True,timeout=180)
            (out/'reference.stdout.log').write_text(ref.stdout)
            (out/'reference.stderr.log').write_text(ref.stderr)
            report['reference_sha256']=sha(SCRIPTS/'wb_reference.py')
            if ref.returncode:
                raise RuntimeError('Reference failed')
            for name in ['app_results.json','truth_16.json','reference_comparison.json']:
                shutil.copy2(data/name,out/name)
            comp=json.loads((out/'reference_comparison.json').read_text())
            ratio_rows=[]
            for image in comp['images']:
                if not image['kind'].startswith('8-bit'):
                    continue
                anchor=image['lanes'][0]
                for lane in image['lanes']:
                    app=lane['appNet']/anchor['appNet']
                    ref=lane['refNet']/anchor['refNet']
                    ratio_rows.append({'image':image['file'],'lane':lane['lane'],'app_ratio':app,'reference_ratio':ref,'absolute_difference':abs(app-ref)})
            ratios={'rows':ratio_rows,'maximum_absolute_difference':max(r['absolute_difference'] for r in ratio_rows)}
            (out/'ratio_comparison.json').write_text(json.dumps(ratios,indent=2))
            with (out/'ratio_comparison.csv').open('w',newline='') as handle:
                writer=csv.DictWriter(handle,fieldnames=['image','lane','app_ratio','reference_ratio','absolute_difference'])
                writer.writeheader()
                writer.writerows(ratio_rows)
            report['real_images']=sum(im['kind'].startswith('8-bit') for im in comp['images'])
            report['real_lanes']=len(ratio_rows)
            report['max_net_relative_difference']=comp['maxNetRelDiff8bit']
            report['max_ratio_absolute_difference']=ratios['maximum_absolute_difference']
            report['status']='passed'
    except Exception as exc:
        report['error']=repr(exc)
    report['finished_utc']=datetime.now(timezone.utc).isoformat()
    (out/'run-report.json').write_text(json.dumps(report,ensure_ascii=False,indent=2))
    print(json.dumps({k:v for k,v in report.items() if k in ['status','error','real_images','real_lanes','max_net_relative_difference','max_ratio_absolute_difference']},indent=2))
    return 0 if report['status']=='passed' else 1

if __name__=='__main__':
    raise SystemExit(main())
```
