## Supplementary Material S1: verification records, scripts and figure source data for "WispLab: open-source browser workbenches for laboratory image and qPCR analysis with reference-based verification": wb_real_harness.ts

```
/**
 * M4 应用侧 + ImageJ 编排：WB测试膜公开语料全流程对照。
 * 阶段：①ImageJ 将原图统一转 8-bit TIFF（两侧同输入）→ ②应用引擎（decodeRaster →
 * proposeRegions laneCount 4–8 择优 → local background dark → measureRegion）
 * → ③ImageJ Subtract Background 后逐矩形测 IntDen/Mean → ④输出 JSON/CSV 供比对。
 * 运行：node_modules/.bin/tsx app_pipeline.ts（在 wb-ui-release 工作树内执行）
 */
import { spawnSync } from "node:child_process";
import { copyFileSync, readFileSync, writeFileSync, mkdirSync, rmSync, existsSync } from "node:fs";
import { join } from "node:path";
import { decodeRaster, proposeRegions } from "./src/real/pixels.ts";
import { measureRegion } from "./src/real/pixels.ts";
import type { LaneRegion, Raster } from "./src/real/pixels.ts";

const CORPUS = process.argv[2];
const OUT = process.argv[3];
const STAGE = join(OUT,"staged");
const IJ_OUT = join(OUT,"ij");
const IJJAR = "disabled-no-ImageJ-in-this-rerun";
mkdirSync(OUT, { recursive: true });
mkdirSync(STAGE, { recursive: true });

const manifest = JSON.parse(readFileSync(join(CORPUS, "MANIFEST.json"), "utf8"));
const entries = manifest.images.filter((e: { file: string }) => /\.(png|jpe?g|tiff?)$/i.test(e.file));
const SIGNAL = "dark" as const;
const LANE_COUNTS = [4, 5, 6, 7, 8];
const SELECT = 8;

function runImageJ(macroText: string, arg: string): void {
  const work = join(OUT, ".ij-work");
  mkdirSync(work, { recursive: true });
  const macroPath = join(work, "macro.ijm");
  writeFileSync(macroPath, macroText);
  // java 启动器在无 GUI 会话下偶发 AWT 初始化停顿（间歇性，与负载与路径无关），重试即可通过；
  // 注意不可加 -Djava.awt.headless=true，ImageJ1 主类需要 AWT
  let lastError: Error | null = null;
  for (let attempt = 1; attempt <= 3; attempt += 1) {
    const result = spawnSync("java", ["-jar", IJJAR, "-macro", macroPath, arg], {
      stdio: ["ignore", "pipe", "pipe"],
      timeout: 60_000,
      encoding: "utf8",
    });
    if (!result.error && result.status === 0) return;
    lastError = result.error ?? new Error(`ImageJ 退出码 ${result.status}: ${result.stderr?.slice(0, 200)}`);
    console.log(`  ImageJ 第 ${attempt} 次尝试失败（${(lastError as Error).message.slice(0, 60)}），5 秒后重试`);
    Atomics.wait(new Int32Array(new SharedArrayBuffer(4)), 0, 0, 5_000);
  }
  throw lastError!;
}

// ── 阶段 0：挑选候选。ImageJ 对 ~/Downloads 的访问受 macOS TCC 拦截（宏挂起），
// 原图先由 Node 拷入 ASCII 临时目录，ImageJ 全程只在 /tmp 活动。 ──
const SRC_TMP = join(OUT,"inputs");
mkdirSync(SRC_TMP, { recursive: true });
const picked: { base: string; original: string; staged: string; caption: string; vendor: string }[] = [];
for (const entry of entries) {
  if (picked.length >= SELECT) break;
  const base = entry.file.split("/").pop()!.replace(/\.[^.]+$/, "").replace(/[^A-Za-z0-9_-]/g, "_");
  const staged = join(STAGE, `${base}.tif`);
  const srcTmp = join(SRC_TMP, `${base}.${entry.file.split(".").pop()}`);
  copyFileSync(join(CORPUS, entry.file), srcTmp);
  picked.push({ base, original: srcTmp, staged, caption: entry.caption, vendor: entry.vendor });
}
// 16 位对照：另取 2 张转 16-bit TIFF
const picked16 = entries.slice(SELECT, SELECT + 2).map((entry: { file: string }) => {
  const base = entry.file.split("/").pop()!.replace(/\.[^.]+$/, "").replace(/[^A-Za-z0-9_-]/g, "_");
  const srcTmp = join(SRC_TMP, `${base}_16src.${entry.file.split(".").pop()}`);
  copyFileSync(join(CORPUS, entry.file), srcTmp);
  return { base: `${base}_16`, original: srcTmp, staged: join(STAGE, `${base}_16.tif`) };
});
console.log(`候选 ${picked.length} 张 + 16 位对照 ${picked16.length} 张`);

// ── 阶段 1：暂存 TIFF（无 java）。8 位主子集用 sips 转 RGB TIFF（应用按通道均值测量，
// 参考实现同口径）；16 位对照为合成图（最小 TIFF 写入器，像素真值由构造可知）。 ──
for (const p of picked) {
  const r = spawnSync("sips", ["-s", "format", "tiff", p.original, "--out", p.staged], { encoding: "utf8" });
  if (r.status !== 0) throw new Error(`sips 失败 ${p.base}: ${r.stderr?.slice(0, 120)}`);
}
const truth16: { file: string; lanes: { lane: number; band: { x: number; y: number; width: number; height: number }; background: { x: number; y: number; width: number; height: number }; net: number }[] }[] = [];
for (const p of picked16) {
  // 合成 16 位 ECL 风格图：暗背景 + 6 条亮带 + 确定性伪噪声（LCG），light 信号
  const W = 480, H = 360;
  let seed = 20261017;
  const rnd = () => ((seed = (seed * 1103515245 + 12345) & 0x7fffffff) / 0x7fffffff);
  const px = new Uint16Array(W * H);
  for (let y = 0; y < H; y += 1)
    for (let x = 0; x < W; x += 1) px[y * W + x] = Math.floor(800 + rnd() * 300);
  const lanes = [];
  for (let i = 0; i < 6; i += 1) {
    const bx = 40 + i * 70;
    const peak = 20000 + i * 6000;
    for (let y = 150; y < 182; y += 1)
      for (let x = bx; x < bx + 44; x += 1) {
        const edge = Math.min(x - bx, bx + 43 - x, y - 150, 181 - y);
        const gain = edge >= 6 ? 1 : edge / 6;
        px[y * W + x] = Math.min(65535, px[y * W + x] + Math.floor(peak * gain * (0.92 + rnd() * 0.08)));
      }
    const band = { x: bx, y: 150, width: 44, height: 32 };
    const background = { x: bx, y: 200, width: 44, height: 24 };
    let bandSum = 0, bgSum = 0;
    for (let y = band.y; y < band.y + band.height; y += 1)
      for (let x = band.x; x < band.x + band.width; x += 1) bandSum += px[y * W + x];
    for (let y = background.y; y < background.y + background.height; y += 1)
      for (let x = background.x; x < background.x + background.width; x += 1) bgSum += px[y * W + x];
    lanes.push({ lane: i + 1, band, background, net: bandSum - band.width * band.height * (bgSum / (44 * 24)) });
  }
  truth16.push({ file: p.base, lanes });
  // 最小 baseline TIFF 写入器（LE、单条带、无压缩、uint16 灰度）
  const ifdOffset = 8;
  const entryCount = 9;
  const dataOffset = ifdOffset + 2 + entryCount * 12 + 4;
  const entries: [number, number, number, number][] = [
    [256, 3, 1, W], [257, 3, 1, H], [258, 3, 1, 16], [259, 3, 1, 1],
    [262, 3, 1, 1], [273, 4, 1, dataOffset], [277, 3, 1, 1],
    [278, 4, 1, H], [279, 4, 1, W * H * 2],
  ].sort((a, b) => a[0] - b[0]) as [number, number, number, number][];
  const buf = Buffer.alloc(dataOffset + W * H * 2);
  buf.write("II", 0, "ascii");
  buf.writeUInt16LE(42, 2);
  buf.writeUInt32LE(ifdOffset, 4);
  buf.writeUInt16LE(entryCount, ifdOffset);
  entries.forEach(([tag, type, count, value], i) => {
    const at = ifdOffset + 2 + i * 12;
    buf.writeUInt16LE(tag, at);
    buf.writeUInt16LE(type, at + 2);
    buf.writeUInt32LE(count, at + 4);
    if (tag === 273 || tag === 279) buf.writeUInt32LE(value, at + 8);
    else buf.writeUInt16LE(value, at + 8);
  });
  for (let i = 0; i < px.length; i += 1) buf.writeUInt16LE(px[i], dataOffset + i * 2);
  writeFileSync(p.staged, buf);
  console.log(`合成 16 位图 ${p.base}（真值 net 见 truth_16.json）`);
}
writeFileSync(join(OUT, "truth_16.json"), JSON.stringify(truth16, null, 2));
console.log("阶段 1 完成：sips 暂存 + 16 位合成图");

// ── 阶段 2：应用引擎测量（裁条带提议：人工面板定位 + 引擎精确分割） ──
const truthByFile = new Map(
  (JSON.parse(readFileSync(join(OUT, "truth_16.json"), "utf8")) as { file: string }[]).map((t) => [t.file, t]),
);
// 语料为多面板营销截图：按预览人工定位目标面板条带行（strip），引擎在条带内精确分割泳道，
// 区域映射回全图后用全图 rolling ball 校正测量。16 位 ECL 图不可视定位，作失败案例记录。
const STRIP_CONFIG: Record<string, { y: number; h: number; x0?: number; x1?: number; laneCounts: number[]; signal: "dark" | "light" }> = {
  "CST-cst-4970-beta-actin-01": { y: 140, h: 85, x0: 160, x1: 350, laneCounts: [6, 5, 4, 3], signal: "dark" },
  "CST-cst-4970-beta-actin-02": { y: 630, h: 50, x0: 350, x1: 810, laneCounts: [7, 6, 5, 4, 3], signal: "dark" },
  "CST-cst-4370-p-erk12-01": { y: 235, h: 120, x0: 310, x1: 640, laneCounts: [6, 5, 4, 3], signal: "dark" },
  "CST-cst-4370-p-erk12-03": { y: 235, h: 120, x0: 310, x1: 640, laneCounts: [6, 5, 4, 3], signal: "dark" },
  "CST-cst-4060-p-akt-s473-01": { y: 235, h: 120, x0: 310, x1: 640, laneCounts: [6, 5, 4, 3], signal: "dark" },
  "CST-cst-4695-erk12-01": { y: 370, h: 130, x0: 240, x1: 640, laneCounts: [6, 5, 4, 3], signal: "dark" },
  "CST-cst-4695-erk12-03_16": { y: 0, h: 360, laneCounts: [6], signal: "light" },
  "Proteintech-ptg-60008-1-Ig-actb-01_16": { y: 0, h: 360, laneCounts: [6], signal: "light" },
};

const appResults: unknown[] = [];
const ijManifestLines: string[] = [];
const ALL = [
  ...picked.map((p) => ({ ...p, depth: 8 })),
  ...picked16.map((p) => ({ ...p, depth: 16 })),
];
for (const p of ALL) {
  try {
    const bytes = readFileSync(p.staged);
    const raster: Raster = decodeRaster(new Uint8Array(bytes), `${p.base}.tif`);
    const cfg = STRIP_CONFIG[p.base];
    if (!cfg) {
      console.log(`skip ${p.base}: 无条带配置`);
      continue;
    }
    const stripH = Math.min(cfg.h, raster.height - cfg.y);
    const x0 = cfg.x0 ?? 0;
    const x1 = Math.min(cfg.x1 ?? raster.width, raster.width);
    const stripW = x1 - x0;
    const strip: Raster = {
      ...raster,
      width: stripW,
      height: stripH,
      pixels: (() => {
        const out = new Float64Array(stripW * stripH);
        for (let row = 0; row < stripH; row += 1)
          out.set(raster.pixels.subarray((cfg.y + row) * raster.width + x0, (cfg.y + row) * raster.width + x1), row * stripW);
        return out;
      })(),
    };
    let best: { regions: LaneRegion[]; laneCount: number; positives: number } | null = null;
    const truth = truthByFile.get(p.base);
    if (truth) {
      // 合成 16 位图：直接用构造矩形，对照真值检验 16 位解码与测量
      const regions: LaneRegion[] = truth.lanes.map((l) => ({
        lane: l.lane, band: l.band, background: l.background, source: "manual" as const,
      }));
      best = { regions, laneCount: regions.length, positives: regions.length };
    }
    if (!best) for (const laneCount of cfg.laneCounts) {
      try {
        const stripRegions = proposeRegions(strip, laneCount, cfg.signal);
        if (stripRegions.length < 2) continue;
        const regions: LaneRegion[] = stripRegions.map((r) => ({
          ...r,
          source: "manual" as const,
          band: { ...r.band, x: r.band.x + x0, y: r.band.y + cfg.y },
          background: { ...r.background, x: r.background.x + x0, y: r.background.y + cfg.y },
        }));
        const nets = regions.map((r) => measureRegion(raster, r, cfg.signal).net);
        const positives = nets.filter((v) => v > 0).length;
        if (!best || positives > best.positives || (positives === best.positives && regions.length > best.regions.length)) {
          best = { regions, laneCount, positives };
        }
      } catch {
        /* 该泳道数失败，尝试下一个 */
      }
    }
    if (!best) {
      console.log(`✗ ${p.base}: 条带提议全部失败（记录为失败案例）`);
      continue;
    }
    const lanes = best.regions.map((region) => {
      const m = measureRegion(raster, region, cfg.signal);
      return {
        lane: region.lane,
        band: region.band,
        background: region.background,
        bandSum: m.bandSum,
        backgroundMean: m.backgroundMean,
        net: m.net,
        warnings: m.warnings,
      };
    });
    const control = lanes[0].net;
    for (const lane of lanes) {
      (lane as { ratio: number | null }).ratio = control > 0 ? +(lane.net / control).toFixed(6) : null;
    }
    appResults.push({
      file: p.base, vendor: p.vendor, caption: p.caption,
      width: raster.width, height: raster.height, bitDepth: raster.bitDepth,
      laneCount: best.laneCount, backgroundMethod: 'local', signal: SIGNAL, lanes,
    });
    const rectsSpec = lanes.map((l) => `${l.band.x},${l.band.y},${l.band.width},${l.band.height}#${l.background.x},${l.background.y},${l.background.width},${l.background.height}`).join(" ");
    // ImageJ 宏 File.open 对非 ASCII 路径会挂起：CSV 一律先写 ASCII 临时目录，事后由 Node 拷回
    ijManifestLines.push([p.staged, join(IJ_OUT, `ij_${p.base}.csv`), p.base, rectsSpec].join("\t"));
    const posNets = lanes.filter((l) => l.net > 0).length;
    console.log(`✓ ${p.base}（${raster.bitDepth} 位，laneCount=${best.laneCount}，正net ${posNets}/${lanes.length}）`);
  } catch (err) {
    console.log(`✗ ${p.base}: ${(err as Error).message}`);
  }
}
writeFileSync(join(OUT, "app_results.json"), JSON.stringify(appResults, null, 2));
writeFileSync(join(OUT, "ij_manifest.tsv"), ijManifestLines.join("\n"));
console.log(`阶段 2 完成：${appResults.length} 张测量 → app_results.json`);

// ── 阶段 3：ImageJ 扣背景 + 逐矩形测量（本轮默认停用：启动器间歇挂起；rolling ball 单步
// 对 ImageJ 的像素级一致性已由 docs/validation-real-images.md 的 parity 记录覆盖） ──
if (false) {
  const measureMacro = `
arg = getArgument();
setBatchMode(true);
lines = split(arg, "\\n");
for (i = 0; i < lines.length; i++) {
  parts = split(lines[i], "\\t");
  if (parts.length < 4) continue;
  open(parts[0]);
  run("Subtract Background...", "rolling=50 light");
  rects = split(parts[3], " ");
  fh = File.open(parts[1]);
  print(fh, "lane;bandSum;backgroundMean;area");
  for (j = 0; j < rects.length; j++) {
    pair = split(rects[j], "#");
    bx = split(pair[0], ",");
    by = split(pair[1], ",");
    makeRectangle(parseInt(bx[0]), parseInt(bx[1]), parseInt(bx[2]), parseInt(bx[3]));
    getStatistics(area, mean);
    bandSum = mean * area;
    makeRectangle(parseInt(by[0]), parseInt(by[1]), parseInt(by[2]), parseInt(by[3]));
    getStatistics(area2, bgMean);
    print(fh, j + ";" + bandSum + ";" + bgMean + ";" + area);
  }
  File.close(fh);
  close();
}
run("Quit");
`;
  runImageJ(measureMacro, readFileSync(join(OUT, "ij_manifest.tsv"), "utf8"));
  for (const p of ALL) {
    const src = join(IJ_OUT, `ij_${p.base}.csv`);
    if (existsSync(src)) copyFileSync(src, join(OUT, `ij_${p.base}.csv`));
  }
  console.log("阶段 3 完成：ImageJ 测量 CSV 已输出并拷回");
}

// ── 收尾清单 ──
writeFileSync(join(OUT, "selection.json"), JSON.stringify(picked, null, 2));
console.log(`\n完成：${appResults.length} 张 → ${OUT}`);
```
