## Supplementary Material S1: verification records, scripts and figure source data for "WispLab: open-source browser workbenches for laboratory image and qPCR analysis with reference-based verification": wb_reference.py

```
#!/usr/bin/env python3
"""M4 独立参考实现（与 M1/M3 同策略，不依赖应用代码）：
- 8 位真实膜：sips→24 位 BMP 解 RGB → 算术通道均值 → 同矩形 bandSum/backgroundMean/net；
- 16 位合成图：与构造真值（truth_16.json）逐值比，检验 16 位解码与测量路径。
对比 app_results.json 的逐泳道 net 与以泳道 1 为对照的 ratio。"""
import sys
import json
import math
import struct
import subprocess
import tempfile
from pathlib import Path

HERE = Path(sys.argv[1]).resolve()

def read_bmp_rgb(path):
    data = Path(path).read_bytes()
    off = struct.unpack_from('<I', data, 10)[0]
    w = struct.unpack_from('<i', data, 18)[0]
    h = struct.unpack_from('<i', data, 22)[0]
    bpp = struct.unpack_from('<H', data, 28)[0]
    assert bpp == 24, f'BPP {bpp}'
    row_size = (w * 3 + 3) & ~3
    top_down = h < 0  # BMP 高度为负表示自上而下存储
    h_abs = abs(h)
    px = [0.0] * (w * h_abs * 3)
    for row in range(h_abs):
        src = off + row * row_size
        dst_row = row if top_down else h_abs - 1 - row
        base = dst_row * w * 3
        for col in range(w):
            i = src + col * 3
            b, g, r = data[i], data[i + 1], data[i + 2]
            j = base + col * 3
            px[j], px[j + 1], px[j + 2] = r, g, b
    return w, h_abs, px

def mean_channel(px):
    n = len(px) // 3
    out = [0.0] * n
    for i in range(n):
        out[i] = (px[i * 3] + px[i * 3 + 1] + px[i * 3 + 2]) / 3.0
    return out

def rect_measure(channel, width, band, bg):
    def sample(rect):
        s = 0.0
        for y in range(rect['y'], rect['y'] + rect['height']):
            base = y * width
            for x in range(rect['x'], rect['x'] + rect['width']):
                s += channel[base + x]
        return s
    band_sum = sample(band)
    area = band['width'] * band['height']
    bg_mean = sample(bg) / (bg['width'] * bg['height'])
    return band_sum, area, bg_mean

def main():
    app = json.loads((HERE / 'app_results.json').read_text())
    truth16 = {t['file']: t for t in json.loads((HERE / 'truth_16.json').read_text())}
    report = []
    max_rel = 0.0
    max16 = 0.0
    for img in app:
        if img['bitDepth'] == 16:
            truth = truth16[img['file']]
            lanes_out = []
            for lane in img['lanes']:
                t = next(l for l in truth['lanes'] if l['lane'] == lane['lane'])
                diff = abs(lane['net'] - t['net'])
                rel = diff / max(abs(t['net']), 1e-12)
                max16 = max(max16, rel)
                lanes_out.append({'lane': lane['lane'], 'appNet': lane['net'],
                                  'truthNet': round(t['net'], 2), 'relDiff': round(rel, 9)})
            report.append({'file': img['file'], 'kind': '16-bit synthetic vs 构造真值', 'lanes': lanes_out})
            continue
        src = HERE / 'staged' / f"{img['file']}.tif"
        with tempfile.NamedTemporaryFile(suffix='.bmp', delete=False) as tmp:
            bmp = tmp.name
        subprocess.run(['sips', '-s', 'format', 'bmp', str(src), '--out', bmp],
                       check=True, capture_output=True)
        w, h, px = read_bmp_rgb(bmp)
        Path(bmp).unlink()
        ch = mean_channel(px)
        lanes_out = []
        for lane in img['lanes']:
            band_sum, area, bg_mean = rect_measure(ch, w, lane['band'], lane['background'])
            ref_net = area * bg_mean - band_sum  # dark 信号：净信号 = 背景均值×面积 − 条带和
            rel = abs(ref_net - lane['net']) / max(abs(lane['net']), abs(ref_net), 1e-12)
            max_rel = max(max_rel, rel)
            lanes_out.append({'lane': lane['lane'], 'appNet': lane['net'],
                              'refNet': ref_net, 'relDiff': round(rel, 9)})
        report.append({'file': img['file'], 'kind': '8-bit real vs 独立参考', 'lanes': lanes_out})
    out = {'maxNetRelDiff8bit': round(max_rel, 9), 'maxNetRelDiff16bit': round(max16, 9), 'images': report}
    (HERE / 'reference_comparison.json').write_text(json.dumps(out, ensure_ascii=False, indent=2))
    for r in report:
        print(f"== {r['file']}（{r['kind']}）")
        for l in r['lanes']:
            ref = l.get('refNet', l.get('truthNet'))
            print(f"  泳道 {l['lane']}: 应用 {l['appNet']:.1f} / 参考 {ref:.1f}（相对差 {l['relDiff']:.2e}）")
    print(f"\n8 位最大相对差 {max_rel:.2e}；16 位对真值最大相对差 {max16:.2e} → reference_comparison.json")

if __name__ == '__main__':
    main()
```
