## Supplementary Material S1: verification records, scripts and figure source data for "WispLab: open-source browser workbenches for laboratory image and qPCR analysis with reference-based verification": wb_synthetic.ts

```
/** Historical synthetic generator reproduced from app_pipeline.ts; fixtures A/B are intentionally identical repetitions, not independent test images. */
import { readFileSync, writeFileSync, mkdirSync } from 'node:fs';
import { join } from 'node:path';
import { decodeRaster, measureRegion } from './src/real/pixels.ts';
const OUT = process.argv[2];
if (!OUT) throw new Error('Usage: tsx wb_synthetic.ts OUTPUT_DIRECTORY');
mkdirSync(OUT, {recursive:true});
const picked16 = ['synthetic_repeat_a', 'synthetic_repeat_b'].map(base => ({base, staged:join(OUT, base+'.tif')}));
const truth16: { file: string; lanes: { lane: number; band: { x: number; y: number; width: number; height: number }; background: { x: number; y: number; width: number; height: number }; net: number }[] }[] = [];
for (const p of picked16) {
  // 合成 16 位 ECL 风格图：暗背景 + 6 条亮带 + 确定性伪噪声（LCG），light 信号
  const W = 480, H = 360;
  let seed = 20261017;
  const rnd = () => ((seed = (seed * 1103515245 + 12345) & 0x7fffffff) / 0x7fffffff);
  const px = new Uint16Array(W * H);
  for (let y = 0; y < H; y += 1)
    for (let x = 0; x < W; x += 1) px[y * W + x] = Math.floor(800 + rnd() * 300);
  const lanes = [];
  for (let i = 0; i < 6; i += 1) {
    const bx = 40 + i * 70;
    const peak = 20000 + i * 6000;
    for (let y = 150; y < 182; y += 1)
      for (let x = bx; x < bx + 44; x += 1) {
        const edge = Math.min(x - bx, bx + 43 - x, y - 150, 181 - y);
        const gain = edge >= 6 ? 1 : edge / 6;
        px[y * W + x] = Math.min(65535, px[y * W + x] + Math.floor(peak * gain * (0.92 + rnd() * 0.08)));
      }
    const band = { x: bx, y: 150, width: 44, height: 32 };
    const background = { x: bx, y: 200, width: 44, height: 24 };
    let bandSum = 0, bgSum = 0;
    for (let y = band.y; y < band.y + band.height; y += 1)
      for (let x = band.x; x < band.x + band.width; x += 1) bandSum += px[y * W + x];
    for (let y = background.y; y < background.y + background.height; y += 1)
      for (let x = background.x; x < background.x + background.width; x += 1) bgSum += px[y * W + x];
    lanes.push({ lane: i + 1, band, background, net: bandSum - band.width * band.height * (bgSum / (44 * 24)) });
  }
  truth16.push({ file: p.base, lanes });
  // 最小 baseline TIFF 写入器（LE、单条带、无压缩、uint16 灰度）
  const ifdOffset = 8;
  const entryCount = 9;
  const dataOffset = ifdOffset + 2 + entryCount * 12 + 4;
  const entries: [number, number, number, number][] = [
    [256, 3, 1, W], [257, 3, 1, H], [258, 3, 1, 16], [259, 3, 1, 1],
    [262, 3, 1, 1], [273, 4, 1, dataOffset], [277, 3, 1, 1],
    [278, 4, 1, H], [279, 4, 1, W * H * 2],
  ].sort((a, b) => a[0] - b[0]) as [number, number, number, number][];
  const buf = Buffer.alloc(dataOffset + W * H * 2);
  buf.write("II", 0, "ascii");
  buf.writeUInt16LE(42, 2);
  buf.writeUInt32LE(ifdOffset, 4);
  buf.writeUInt16LE(entryCount, ifdOffset);
  entries.forEach(([tag, type, count, value], i) => {
    const at = ifdOffset + 2 + i * 12;
    buf.writeUInt16LE(tag, at);
    buf.writeUInt16LE(type, at + 2);
    buf.writeUInt32LE(count, at + 4);
    if (tag === 273 || tag === 279) buf.writeUInt32LE(value, at + 8);
    else buf.writeUInt16LE(value, at + 8);
  });
  for (let i = 0; i < px.length; i += 1) buf.writeUInt16LE(px[i], dataOffset + i * 2);
  writeFileSync(p.staged, buf);
  console.log(`合成 16 位图 ${p.base}（真值 net 见 truth_16.json）`);
}
writeFileSync(join(OUT, "truth_16.json"), JSON.stringify(truth16, null, 2));

const measurements=[];
for (const [i,p] of picked16.entries()) {
 const raster=decodeRaster(new Uint8Array(readFileSync(p.staged)), p.staged);
 const lanes=truth16[i].lanes.map(l => ({...l, ...measureRegion(raster, {lane:l.lane, band:l.band, background:l.background, source:'manual'}, 'light')}));
 measurements.push({file:p.base, width:raster.width, height:raster.height, bitDepth:raster.bitDepth, lanes});
}
writeFileSync(join(OUT,'app_results.json'),JSON.stringify(measurements,null,2));
console.log('Recomputed TIFF decode + local-background measurement for 12 repeated lane measurements (6 unique constructed lanes).');
```
