## Supplementary Material S1: verification records, scripts and figure source data for "WispLab: open-source browser workbenches for laboratory image and qPCR analysis with reference-based verification": README.md

# Auto-qPCR example-data provenance and license

The qPCR input in this supplement originates from
[`neuroeddu/Auto-qPCR`](https://github.com/neuroeddu/Auto-qPCR), revision
`d881ebd01667766882f90de850bfc26d93c54e27` (4 August 2022), at
[`Example_Data/RELATIVE_example.csv`](https://github.com/neuroeddu/Auto-qPCR/blob/d881ebd01667766882f90de850bfc26d93c54e27/Example_Data/RELATIVE_example.csv).
The upstream CSV was fetched at that fixed revision and was byte-identical to
the fixture in the WispLab qPCR paper revision `4142b6078251d3973667bb59352a016ac0ebf3da`.
Both files have SHA256
`21083876c4b827dea58fd777acb5019569153142c4a9acde5fd852feff6dd8b2`.

The verified upstream license is **GNU General Public License version 3**, not
MIT. The complete, unchanged upstream `LICENCE.txt`, including its copyright
notice, is supplied as `Auto-qPCR-LICENCE.txt`. Its Git blob was verified against
`6eb09dfe91b7e958295cd556f7d871e9b2747259`. This corrects the MIT attribution in
older local provenance notes; those notes are not used as the licensing source.

Upstream copyright notice: Copyright (C) 2021 Rhalena Thomas, Eddie Cai, Gracia Gu
and Iva Demirova at the NeuroEDDU.

The corresponding data-derived materials are
`fresh-20261018/pcr/matrix.json` (CSV parsed to nonempty string rows), and the
`inputRows` portions of the three real-data `app_output_*.json` files. These
reformatted data are accompanied by the upstream GPL-3 license and attribution;
the synthetic S4 Cq example was constructed separately. This notice does not
relicense the WispLab application repositories or unrelated third-party inputs.
No Auto-qPCR application source code or RDML fixtures are copied into this
supplement.

The authoritative upstream license is
[`LICENCE.txt` at the same revision](https://github.com/neuroeddu/Auto-qPCR/blob/d881ebd01667766882f90de850bfc26d93c54e27/LICENCE.txt).
Machine-readable provenance and checksums are in `Auto-qPCR-provenance.json`.

Please also cite Maussion G et al. Auto-qPCR; a python-based web app for automated
and reproducible analysis of qPCR data. Scientific Reports 11, 21293 (2021).
https://doi.org/10.1038/s41598-021-99727-6
