## Supplementary Material S1: verification records, scripts and figure source data for "WispLab: open-source browser workbenches for laboratory image and qPCR analysis with reference-based verification": plot_figures.py

```
#!/usr/bin/env python3
"""Plot WispLab verification records; does not execute the products.

Requires Python 3, numpy and matplotlib. Run from any directory:
    python3 manuscript/scripts/plot_figures.py
The default source snapshots are preserved byte-for-byte beside the figures.
"""
from __future__ import annotations

import argparse
import csv
import hashlib
import json
from pathlib import Path

import matplotlib
matplotlib.use("Agg")
import matplotlib.pyplot as plt
from matplotlib.patches import FancyArrowPatch, FancyBboxPatch
from matplotlib.ticker import ScalarFormatter
import numpy as np

HERE = Path(__file__).resolve().parent.parent
INDIGO, BLUE, ORANGE, PURPLE = "#4F46A5", "#237FA4", "#D78227", "#916092"
INK, MUTED, GRID = "#222A3A", "#5A6475", "#E4E7EF"
COLORS = [INDIGO, BLUE, ORANGE, PURPLE]
MARKERS = ["o", "s", "^", "D"]


def configure():
    plt.rcParams.update({
        "font.family": "DejaVu Sans", "font.size": 9,
        "axes.labelsize": 9, "axes.titlesize": 10,
        "axes.titleweight": "bold", "axes.titlepad": 12,
        "axes.labelcolor": INK, "text.color": INK,
        "xtick.color": MUTED, "ytick.color": MUTED,
        "axes.edgecolor": "#A4ACBB", "axes.spines.top": False,
        "axes.spines.right": False, "axes.axisbelow": True,
        "grid.color": GRID, "grid.linewidth": .6,
        "legend.frameon": False, "legend.fontsize": 8,
        "pdf.fonttype": 42, "ps.fonttype": 42, "svg.fonttype": "none",
        "savefig.dpi": 300, "figure.facecolor": "white",
        "savefig.facecolor": "white",
    })


def write_csv(path, rows):
    if not rows:
        raise ValueError(f"No data to write: {path}")
    with path.open("w", newline="", encoding="utf-8") as f:
        writer = csv.DictWriter(f, fieldnames=list(rows[0]))
        writer.writeheader()
        writer.writerows(rows)


def save(fig, out, name):
    for ext in ("pdf", "svg", "png"):
        fig.savefig(out / f"{name}.{ext}", dpi=300)
    plt.close(fig)


def panel(ax, letter, title):
    ax.set_title(title, loc="left")
    ax.text(-.16, 1.075, letter, transform=ax.transAxes,
            fontsize=13, fontweight="bold", va="bottom")
    ax.grid(True)


def equality(ax, low, high, log=False):
    ax.plot([low, high], [low, high], "--", color=MUTED, lw=.9, zorder=0)
    ax.set_xlim(low, high)
    ax.set_ylim(low, high)
    if log:
        ax.set_xscale("log")
        ax.set_yscale("log")


def figure_overview(out):
    fig, ax = plt.subplots(figsize=(10.8, 6.5))
    ax.set_xlim(0, 10.8)
    ax.set_ylim(0, 6.5)
    ax.axis("off")

    def box(x, y, w, h, title, body, fill="#F3F2FC", border=INDIGO):
        ax.add_patch(FancyBboxPatch((x, y), w, h,
            boxstyle="round,pad=.035,rounding_size=.09", fc=fill,
            ec=border, lw=.85))
        ax.text(x + w/2, y+h-.23, title, ha="center", va="top",
                fontsize=10, weight="bold")
        ax.text(x + w/2, y+h-.63, body, ha="center", va="top",
                fontsize=8.6, linespacing=1.55, color=MUTED)

    def arrow(start, end, dashed=False, color=MUTED):
        ax.add_patch(FancyArrowPatch(start, end, arrowstyle="-|>",
            mutation_scale=11, lw=1, color=color,
            linestyle="--" if dashed else "-"))

    ax.text(.4, 6.18, "WispLab: default browser workflow and verification scope",
            fontsize=13, weight="bold")
    ax.text(.4, 5.82, "Three workbenches share a web entry point; assay calculations execute in the browser.",
            fontsize=9, color=MUTED)
    box(.45, 3.8, 2.9, 1.65, "Western blot", "TIFF / PNG → lane regions\nLocal background → net signal\nLane-1 normalization")
    box(3.95, 3.8, 2.9, 1.65, "Microscopy", "IHC / stains / fluorescence\nRegion selection → image analysis\nMasks and quantitative summaries")
    box(7.45, 3.8, 2.9, 1.65, "qPCR", "Cq tables → replicate aggregation\nReference genes → relative expression\nStatistical comparisons")
    for x in (1.9, 5.4, 8.9):
        arrow((x, 3.76), (x, 3.34))
    box(.45, 2.3, 9.9, 1.0, "Review and export", "Inspect regions, settings and results; save tables, figures and analysis records.", fill="#FAFBFD", border="#A4ACBB")
    arrow((5.4, 2.25), (5.4, 1.87))
    box(.45, .53, 9.9, 1.3, "Recorded numerical checks", "WB: 4 public membranes / 21 lanes   •   IHC area: 8 images / 10 regions\nqPCR: 60-well example + synthetic scenario   •   Independent Python references", fill="#F3F6FC", border=BLUE)
    ax.text(.48, .11, "Optional external AI services are outside this default local flow and may transmit user-selected data.",
            fontsize=8, color=MUTED)
    fig.subplots_adjust(left=0, right=1, top=1, bottom=0)
    save(fig, out, "figure1_workflow")


def figure_wb(data, synthetic_data, out):
    rows, synthetic = [], []
    short = {
        "CST-cst-4970-beta-actin-01": "β-actin · 4970",
        "CST-cst-4370-p-erk12-01": "p-ERK1/2 · 4370",
        "CST-cst-4060-p-akt-s473-01": "p-AKT · 4060",
        "CST-cst-4695-erk12-01": "ERK1/2 · 4695",
    }
    for record in data["images"]:
        for lane in record["lanes"]:
            if "refNet" in lane:
                a, r = lane["appNet"], lane["refNet"]
                rows.append({"image_id": record["file"], "lane": lane["lane"],
                    "app_net": a, "reference_net": r,
                    "absolute_difference_from_serialized_values": abs(a-r),
                    "relative_difference_as_recorded": lane["relDiff"]})
    for lane in synthetic_data["measurements"]:
        synthetic.append({"synthetic_image_id": lane["file"], "lane": lane["lane"],
            "app_net": lane["app_net"], "truth_net": lane["constructed_net"],
            "absolute_difference_as_recorded": lane["absolute_difference"]})
    assert len(rows) == 21 and len({r["image_id"] for r in rows}) == 4
    assert len(synthetic) == 12
    write_csv(out / "figure2_wb_source.csv", rows)
    write_csv(out / "wb_synthetic_16bit_source.csv", synthetic)
    fig, axes = plt.subplots(1, 2, figsize=(10.8, 4.7))
    offset = 0
    for i, (key, label) in enumerate(short.items()):
        group = [r for r in rows if r["image_id"] == key]
        x = np.array([r["reference_net"] for r in group])
        y = np.array([r["app_net"] for r in group])
        axes[0].scatter(x, y, s=38, color=COLORS[i], marker=MARKERS[i],
                        label=f"{label} (n = {len(group)})", alpha=.85)
        axes[1].scatter(np.arange(offset+1, offset+len(group)+1), abs(y-x),
                        s=38, color=COLORS[i], marker=MARKERS[i], alpha=.85)
        offset += len(group)
    equality(axes[0], 0, 450000)
    axes[0].set_xlabel("Reference net signal (gray-level × pixel)")
    axes[0].set_ylabel("WispLab net signal (gray-level × pixel)")
    axes[0].ticklabel_format(axis="both", style="sci", scilimits=(0, 0), useMathText=True)
    axes[0].legend(loc="upper left", handlelength=1.0, labelspacing=.55)
    axes[1].set_xlabel("Lane measurement index (source-table order)")
    axes[1].set_ylabel("Absolute net difference (gray-level × pixel)")
    axes[1].set_xlim(.5, 21.5)
    axes[1].set_xticks(range(1, 22))
    axes[1].tick_params(axis="x", labelsize=7)
    maximum_error = max(r["absolute_difference_from_serialized_values"] for r in rows)
    if maximum_error == 0:
        axes[1].set_ylim(-.1, .1)
        axes[1].set_yticks([0])
        axes[1].text(.5, .84, "21 / 21 net signals identical\nMaximum absolute difference = 0",
                     transform=axes[1].transAxes, va="top", ha="center", fontsize=9)
    else:
        axes[1].set_ylim(-maximum_error*.04, maximum_error*1.2)
    axes[1].axhline(0, lw=.7, color=MUTED, zorder=0)
    panel(axes[0], "A", "Net-signal agreement")
    panel(axes[1], "B", "Magnitude of numerical differences")
    fig.text(.08, .025, "Fixed-tag rerun: 4 public membranes · 21 lanes · local background · shared TIFF inputs and matched rectangles", fontsize=8, color=MUTED)
    fig.subplots_adjust(left=.08, right=.98, bottom=.19, top=.87, wspace=.32)
    save(fig, out, "figure2_wb_agreement")
    return {"public_images": 4, "lanes": len(rows),
        "max_relative_net_difference_as_recorded": max(r["relative_difference_as_recorded"] for r in rows),
        "max_absolute_net_difference_from_serialized_values": max(r["absolute_difference_from_serialized_values"] for r in rows),
        "synthetic_16bit_files": 2, "synthetic_recorded_lane_measurements": len(synthetic),
        "synthetic_unique_constructed_lanes": 6,
        "synthetic_replication_note": "The two files repeat the same constructed six-lane pattern and are not independent images.",
        "synthetic_max_absolute_difference_as_recorded": max(r["absolute_difference_as_recorded"] for r in synthetic)}


def figure_ihc(data, out):
    rows = []
    for i, r in enumerate(data["rows"], start=1):
        rows.append({"roi_id": f"R{i:02d}", "image_id": r["file"],
            "roi_as_recorded": r["roi"], "reference_pixel_count": r["n"],
            "app_positive_area_percent": r["appPct"],
            "reference_positive_area_percent": r["referencePct"],
            "reference_area_at_app_threshold_percent": r["referencePctAtAppThr"],
            "absolute_difference_independent_thresholds_pp_as_recorded": r["absDiffPct"],
            "absolute_difference_same_threshold_pp_as_recorded": r["absDiffAtSameThr"],
            "reference_threshold": r["referenceThreshold"], "app_threshold": r["appThreshold"]})
        rows[-1].update({"roi_rectangle_xywh": json.dumps(r.get("roi_rectangle_xywh")),
            "application_roi_name": r.get("application_roi_name", r["roi"]),
            "mapping_corrected_on_20261018": r.get("mapping_changed", False),
            "historical_app_pct_before_rematching": r.get("original_appPct", r["appPct"])})
    assert len(rows) == 10 and len({r["image_id"] for r in rows}) == 8
    write_csv(out / "figure3_ihc_source.csv", rows)
    x = np.array([r["reference_positive_area_percent"] for r in rows])
    y = np.array([r["app_positive_area_percent"] for r in rows])
    auto = np.array([r["absolute_difference_independent_thresholds_pp_as_recorded"] for r in rows])
    same = np.array([r["absolute_difference_same_threshold_pp_as_recorded"] for r in rows])
    fig, axes = plt.subplots(1, 2, figsize=(10.8, 4.7))
    equality(axes[0], 0, 46)
    axes[0].scatter(x, y, s=45, color=INDIGO, alpha=.85)
    for i in (3, 6, 7, 9):
        axes[0].annotate(rows[i]["roi_id"], (x[i], y[i]), xytext=(4, 6), textcoords="offset points", fontsize=7.5)
    axes[0].set_xlabel("Reference positive area (%)")
    axes[0].set_ylabel("WispLab positive area (%)")
    axes[0].text(.05, .93, f"8 images · 10 regions\nMax difference: {max(auto):.4f} percentage points", transform=axes[0].transAxes, va="top", color=MUTED, fontsize=8.3)
    yy = np.arange(len(rows))
    axes[1].hlines(yy, np.minimum(auto, same), np.maximum(auto, same), color="#B4BAC9", lw=1)
    axes[1].scatter(auto, yy, s=35, color=INDIGO, marker="o", label="Independent Otsu thresholds", zorder=3)
    axes[1].scatter(same, yy, s=35, color=ORANGE, marker="D", facecolors="none", label="Reference at app threshold", zorder=3)
    axes[1].set_yticks(yy, [r["roi_id"] for r in rows])
    axes[1].invert_yaxis()
    max_diff = max(max(auto), max(same))
    axes[1].set_xlim(-max_diff*.035, max_diff*1.15)
    axes[1].set_xlabel("Absolute area difference (percentage points)")
    axes[1].legend(loc="upper right", fontsize=7.5, handlelength=1.1)
    axes[1].set_title("Area differences by region", loc="left", pad=12)
    panel(axes[0], "A", "Positive-area comparison")
    axes[1].text(-.16, 1.075, "B", transform=axes[1].transAxes, fontsize=13, weight="bold", va="bottom")
    axes[1].grid(axis="x")
    fig.text(.08, .025, "Matched preprocessing and ROI geometry; fixed-tag application and independent reference reruns (18 Oct 2026).\nOtsu thresholds matched in all images; paired markers overlap. Reference code is not annotated ground truth.",
             fontsize=7.7, color=MUTED)
    fig.subplots_adjust(left=.08, right=.98, bottom=.18, top=.87, wspace=.32)
    save(fig, out, "figure3_ihc_area")
    return {"images": 8, "regions": len(rows),
        "audit_type": data.get("audit_type"),
        "application_engine_sha": data.get("application_engine_sha"),
        "max_absolute_difference_independent_thresholds_pp_as_recorded": float(max(auto)),
        "max_absolute_difference_same_threshold_pp_as_recorded": float(max(same)),
        "median_absolute_difference_independent_thresholds_pp_as_recorded": float(np.median(auto)),
        "count_independent_threshold_difference_below_0_6_pp": int(sum(auto < .6)),
        "scope": "DAB/IHC area only; not H&E, trichrome, IF, cell-count or H-score validation"}


def figure_pcr(data, out):
    folds, dunnett, checks = [], [], []
    for scenario in data["scenarios"]:
        s = scenario["name"]
        checks.append({"scenario": s, "passed": scenario["passed"], "failed": scenario["failed"], "all_pass": scenario["all_pass"]})
        for c in scenario["checks"]:
            if c["field"].endswith(".foldChange") and not s.startswith("S4"):
                folds.append({"scenario": s, "field": c["field"], "app": c["app"], "reference": c["ref"],
                              "absolute_difference": abs(c["app"]-c["ref"])})
            if c["field"].endswith(".dunnettP"):
                dunnett.append({"scenario": s, "field": c["field"], "app": c["app"], "reference": c["ref_quad"],
                    "absolute_difference": abs(c["app"]-c["ref_quad"]), "relative_difference": c["rel_diff"],
                    "raw_p_lower_bound": c["raw_p_interval"][0], "bonferroni_upper_bound": c["raw_p_interval"][1],
                    "plotted": not s.startswith("S3"),
                    "plot_note": "S3 repeats the same statistical contrasts as S2; not plotted twice" if s.startswith("S3") else ""})
    unique = [r for r in dunnett if r["plotted"]]
    assert len(unique) == 9 and len(folds) == 32
    assert sum(r["failed"] for r in checks) == 0
    write_csv(out / "figure4_pcr_fold_change_source.csv", folds)
    write_csv(out / "figure4_pcr_dunnett_source.csv", dunnett)
    write_csv(out / "pcr_check_counts.csv", checks)
    fig, axes = plt.subplots(1, 3, figsize=(12.1, 4.6), gridspec_kw={"width_ratios": [1, 1, 1.08]})
    names = {"S1_livak_welch": "S1: Livak (n = 8)", "S2_livak_anova": "S2: Livak (n = 12)", "S3_pfaffl": "S3: Pfaffl (n = 12)"}
    for i, (s, name) in enumerate(names.items()):
        group = [r for r in folds if r["scenario"] == s]
        axes[0].scatter([r["reference"] for r in group], [r["app"] for r in group], s=36,
                        marker=MARKERS[i], facecolors="none", edgecolors=COLORS[i], label=name)
    low = min(r["reference"] for r in folds)*.72
    high = max(r["reference"] for r in folds)*1.3
    equality(axes[0], low, high, log=True)
    axes[0].set_xlabel("Reference fold change")
    axes[0].set_ylabel("WispLab fold change")
    axes[0].legend(loc="upper left", fontsize=7.5)
    axes[0].text(.97, .06, "32 outputs\nSome points overlap", transform=axes[0].transAxes,
                 fontsize=7.5, color=MUTED, ha="right")
    for s, color, marker, label in (("S2_livak_anova", INDIGO, "o", "S2: public Cq example (n = 6)"), ("S4_moderate_t", ORANGE, "^", "S4: synthetic Cq (n = 3)")):
        group = [r for r in unique if r["scenario"] == s]
        axes[1].scatter([r["reference"] for r in group], [r["app"] for r in group],
                        s=40, color=color, marker=marker, label=label)
    equality(axes[1], 4e-7, .05, log=True)
    axes[1].set_xlabel("Reference adjusted p")
    axes[1].set_ylabel("WispLab adjusted p")
    axes[1].legend(loc="upper left", fontsize=7)
    for i, r in enumerate(unique):
        color = INDIGO if r["scenario"].startswith("S2") else ORANGE
        marker = "o" if r["scenario"].startswith("S2") else "^"
        axes[2].scatter(r["absolute_difference"], i, s=35, color=color, marker=marker)
    labels = []
    for r in unique:
        name = r["field"].replace(".dunnettP", "").replace("AiW002-2-D7", "AiW D7").replace("KYOU-D7", "KYOU D7")
        labels.append(name)
    axes[2].set_yticks(np.arange(len(unique)), labels, fontsize=7)
    axes[2].invert_yaxis()
    axes[2].set_xscale("log")
    axes[2].set_xlim(3e-12, 2e-7)
    axes[2].set_xlabel("Absolute adjusted-p difference")
    for ax, letter, title in zip(axes, "ABC", ["Relative-expression agreement", "Dunnett agreement after correction", "Adjusted-p differences"]):
        panel(ax, letter, title)
    axes[2].text(.05, .04, f"Max relative difference\n{max(r['relative_difference'] for r in unique):.3g}", transform=axes[2].transAxes, fontsize=7.5, color=MUTED)
    fig.text(.06, .035, "Fixed-tag rerun: 4 scenarios, 422 checks passed. S3 Dunnett values repeat S2 and are excluded from panels B–C.",
             fontsize=8, color=MUTED)
    fig.subplots_adjust(left=.06, right=.985, bottom=.19, top=.86, wspace=.62)
    save(fig, out, "figure4_pcr_agreement")
    return {"scenarios": len(checks), "recorded_passed_checks": sum(r["passed"] for r in checks),
        "recorded_failed_checks": sum(r["failed"] for r in checks),
        "fold_change_outputs_plotted": len(folds),
        "fold_change_max_absolute_difference": max(r["absolute_difference"] for r in folds),
        "unique_dunnett_comparisons_plotted": len(unique), "public_data_dunnett_comparisons": 6,
        "synthetic_dunnett_comparisons": 3,
        "dunnett_max_relative_difference": max(r["relative_difference"] for r in unique),
        "dunnett_max_absolute_difference": max(r["absolute_difference"] for r in unique)}


def main():
    parser = argparse.ArgumentParser(description=__doc__)
    parser.add_argument("--source-dir", type=Path, default=HERE / "figures/source-data")
    parser.add_argument("--output-dir", type=Path, default=HERE / "figures")
    args = parser.parse_args()
    args.output_dir.mkdir(parents=True, exist_ok=True)
    sources = {}
    hashes = {}
    for key, filename in (("wb", "wb_reference_comparison_pinned_20261018.json"), ("wb_synthetic", "wb_synthetic_comparison_pinned_20261018.json"), ("ihc", "ihc_reference_area_comparison_pinned_20261018.json"), ("pcr", "pcr_comparison_report_pinned_20261018.json")):
        p = args.source_dir / filename
        blob = p.read_bytes()
        sources[key] = json.loads(blob)
        hashes[filename] = hashlib.sha256(blob).hexdigest()
    configure()
    figure_overview(args.output_dir)
    summary = {"purpose": "All numerical figures use fixed-tag application and independent-reference reruns on 2026-10-18 with existing dependencies reused. WB sides share freshly converted TIFF inputs; IHC preprocessing and ROI geometry were aligned. No new biological experiment or clean dependency-installation test is claimed.",
        "source_sha256": hashes, "wb": figure_wb(sources["wb"], sources["wb_synthetic"], args.output_dir),
        "ihc": figure_ihc(sources["ihc"], args.output_dir),
        "pcr": figure_pcr(sources["pcr"], args.output_dir),
        "plotting_environment": {"matplotlib": matplotlib.__version__, "numpy": np.__version__}}
    (args.output_dir / "figure-data-summary.json").write_text(json.dumps(summary, indent=2)+"\n", encoding="utf-8")
    print(json.dumps(summary, indent=2))


if __name__ == "__main__":
    main()
```
